# Streamlined Production of Recombinant Adeno-Associated Viruses Using Gateway Cloning Technology for Neural Circuit All-Optical Interrogation

**DOI:** 10.64898/2026.09.22.752944

**Authors:** Annick Ayon, Abdelali Jalil, Coailinn Egan, Bertrand Ducos, Caroline Mailhes-Hamon, Astou Tangara, Benjamin Mathieu, Alexis-Pierre Bemelmans, Isabel Llano, Laurent Bourdieu, Stéphane Dieudonné, Vincent Villette, Jonathan Bradley

## Abstract

The advent of optogenetics and the rapid development of genetically encoded actuators and reporters for calcium, voltage, neurotransmitters, and other molecules have revolutionized neuroscience research by enabling precise, non-invasive optical interrogation of neural circuits. Successful implementation, however, critically depends on efficient reporter expression in defined neuronal populations. We present a Gateway®-based cloning platform providing a robust pipeline for rapid, efficient construction of recombinant adeno-associated virus (rAAV) vectors optimized for neuroscience applications. This platform addresses promoter selection, indicator engineering, and capsid-type optimization, and enables systematic optimization of vector components across a range of optical tools. The benefit of this modular approach is illustrated in the case of ULoVE, a two-photon microscopy method based on acousto-optic deflectors (AODs). This method provides serial light-targeting with kHz sampling rates and high signal-to-noise ratio in vivo, which imposes stringent requirements on indicator expression — including cell-type specificity, sparse labelling, and precise control of expression levels— efficiently met through the combinatorial flexibility of the Gateway pipeline. As specific examples, Gateway-constructed Cre-driver viruses combined with Cre-dependent reporters enabled cell-type-specific labelling for two complementary applications. In cerebellar Purkinje cells, an L7::Cre driver virus paired with a Gateway-constructed voltage indicator (JEDI2P-Kv) and ULoVE two-photon excitation at ∼5 kHz enables resolution of sub-millisecond dendritic voltage dynamics in awake, behaving mice, including optical detection of dendritic spikelets previously accessible only via intracellular electrophysiology. The same driver virus paired with calcium indicators (GCaMP6f, jRGECO1a) resolves climbing fiber-evoked calcium kinetics under the same conditions. In acute cerebellar slices, a kit::Cre driver virus enabled cell-type-specific expression of the optogenetic actuator ChR2(H134R) in molecular layer interneurons, where ULoVE doughnut-pattern photostimulation achieved single-cell, sub-millisecond optogenetic activation with micron-scale spatial resolution.

Beyond ULoVE applications, the same sparse, strong labelling also proved suitable for anatomical tracing of axonal projections in cleared cerebellar tissue, resolving individual deep cerebellar nuclei (DCN) axon terminals at the granule cell layer–molecular layer boundary. Taken together, our dual-front approach—combining a modular AAV expression pipeline with AOD-based optical acquisition—provides an integrated platform for developing sophisticated optical tools, including but not limited to AOD-based microscopy, for neural circuit interrogation.

## Introduction

Understanding neural circuit function requires tools capable of monitoring and manipulating neuronal activity with high spatiotemporal precision. The development of optogenetic actuators and genetically encoded indicators of calcium (GECIs), voltage (GEVIs), neurotransmitters (e.g., glutamate and glycine), and neuromodulators (e.g., dopamine) has transformed neuroscience by enabling researchers to manipulate and observe neuronal activity in behaving animals (Marvin et al., 2013; Deisseroth, 2015; Lin and Schnitzer, 2016; Patriarchi et al., 2018; Sun et al., 2018; Zhang et al., 2018). Channelrhodopsin-2, the founding optogenetic tool, permits millisecond-scale control of neuronal firing in response to blue light stimulation (Boyden et al., 2005). Complementary to optogenetic activation, GECIs such as GCaMP enable real-time visualization of neural activity by reporting intracellular calcium fluctuations through fluorescence changes (Chen et al., 2013). Unlike GECIs, which monitor calcium as a proxy for neuronal activity, GEVIs directly report voltage fluctuations through fluorescence changes (Knöpfel and Song, 2019).

The practical implementation of these technologies requires efficient gene delivery systems capable of achieving robust, cell-type-specific expression in neural tissue. Recombinant adeno-associated viruses (rAAVs) have emerged as the predominant vector platform for neuroscience applications because of their excellent safety profile, low immunogenicity, and ability to drive long-term transgene expression in neurons (Deverman et al., 2016; Chan et al., 2017). It is important to note, however, that rAAVs are not entirely inert in neural tissue: recent work has shown that an innate immune response to the unmethylated CpG-rich AAV genome, mediated by Toll-like receptor 9 (TLR9), can reduce dendritic complexity and disrupt synaptic transmission in cortical neurons at experimentally relevant titers, independent of capsid type, transgene, or promoter (Suriano et al., 2024). These effects, which can be mitigated by CpG depletion or TLR9 blockade, are a relevant consideration for studies using rAAVs to interrogate neuronal structure and function, and underscore the value of expression strategies — such as sparse, titrated labelling — that minimize the viral genome load delivered to any individual cell (Guo et al., 2023). Optimizing expression in this way, however, is only practical if the underlying vectors can be built and modified rapidly. The iterative nature of neuroscience research — continually refining promoters, coding sequences, regulatory elements, and expression strategies to balance signal quality against viral load and off-target effects — therefore demands cloning strategies that are flexible, robust, and efficient.

Gateway® cloning technology, based on bacteriophage λ site-specific recombination, addresses key limitations of conventional restriction enzyme-based cloning, namely its dependence on suitable unique restriction sites, variable ligation efficiency, and the need to re-engineer constructs whenever a component is exchanged, by providing a modular, high-efficiency system for DNA manipulation (Landy, 1989; Hartley et al., 2000). During lambda phage infection of bacterial cells, recombination events occur through the interaction of phage attachment sequences (attP) with bacterial attachment sequences (attB). When attP and attB sequences recombine, the phage genome inserts itself into the bacterial chromosome (lysogenic phase), creating two novel recombination sequences on either side of the inserted genome (attL on the left and attR on the right), as shown in **Figure 1**.

**Figure 1.**
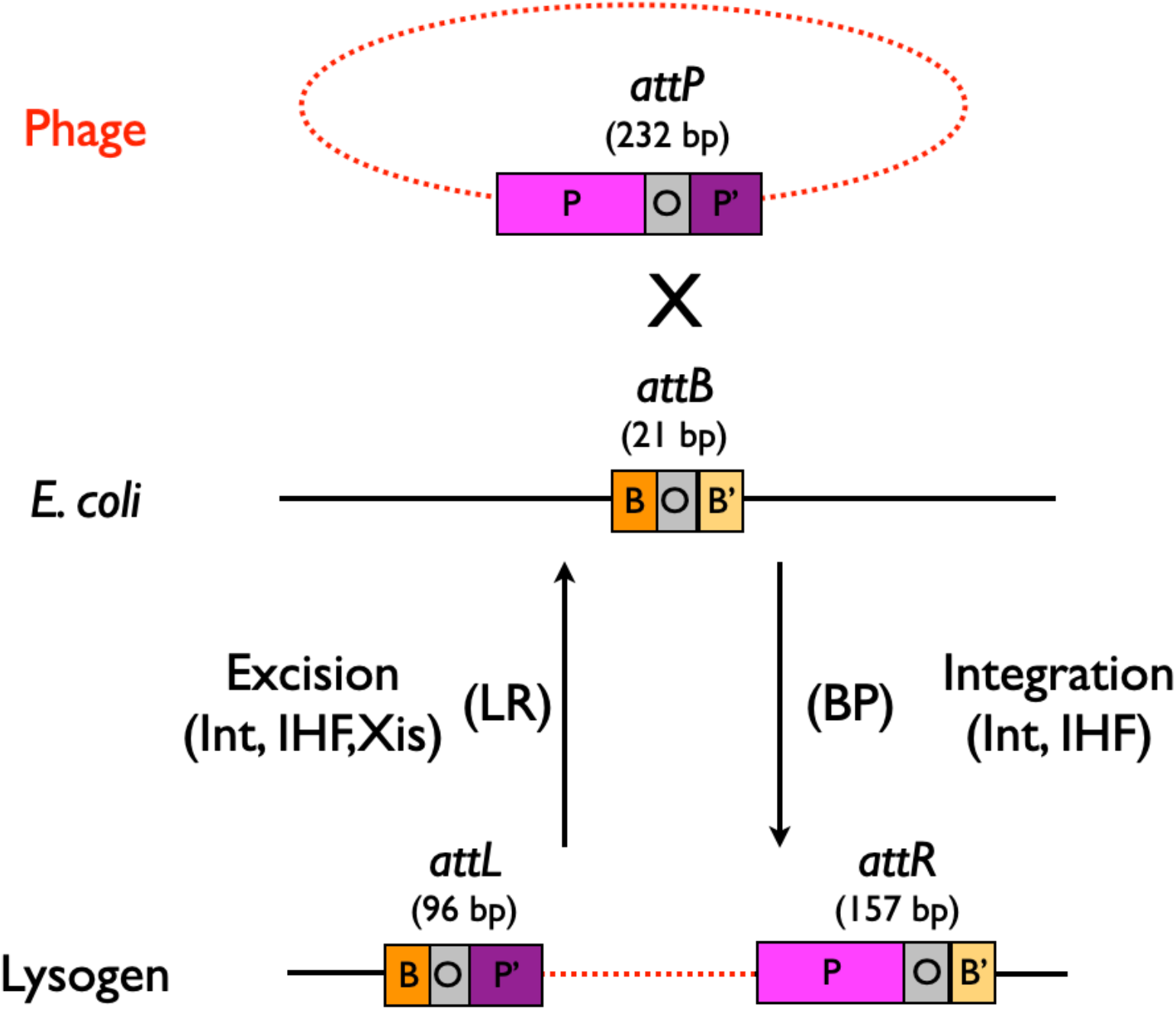
Site-specific recombination in phage λ. The integrative and excisive λ recombination pathways are shown. BP stands for ‘attB x attP recombination’, whereas LR stands for ‘attL x attR recombination’. The attP and attB sites are each composed of a common 7-base pair nucleotide sequence referred to as the core sequence (O, grey) and two flanking arms called P and P′ (light and dark pink), and B and B′ (dark and light orange), respectively. Recombination begins with phage-coded integrase (Int) and bacterial integration host factor (IHF) binding tightly to the 232-base pair attP. The complex then couples with the 21-base pair attB in the bacterial chromosome. Staggered DNA nicks are produced at the ends of the core sequence of both attP and attB sequences, followed by strand exchange between them, with the O sequence providing homology between the two joined sequences, allowing a tiny heteroduplex joint to form at this point of exchange. For the integration event, only Int and IHF are involved, whereas excision also requires the excisionase (Xis), an additional phage-encoded protein.

Under specific circumstances, attL and attR sequences can undergo recombination, causing the phage to be removed from the bacterial DNA while restoring the original attP and attB sequences (lytic phase). Gateway vectors utilize engineered versions of these attachment sequences, enabling efficient insertion of target DNA fragments. The Gateway system operates through two fundamental reactions **(Figure 2, Supplement to Figure 2)**: BP Clonase enzymes catalyze recombination between attB-flanked DNA fragments of interest and attP-containing donor plasmids (pDONR), producing entry clones (pENTR) containing the desired sequence bracketed by attL sites; LR Clonase enzymes then mediate recombination between pENTR attL sites and the cognate attR sites of a destination plasmid (pDEST), producing a functional expression clone (pEXPR) while simultaneously excising a ccdB negative-selection cassette from it. The modularity of this system — in which any promoter entry clone can be combined with any transgene entry clone in a single 1.5-hour reaction, then plated for colony growth overnight— makes it ideally suited to the combinatorial demands of optical neuroscience.

**Figure 2.**
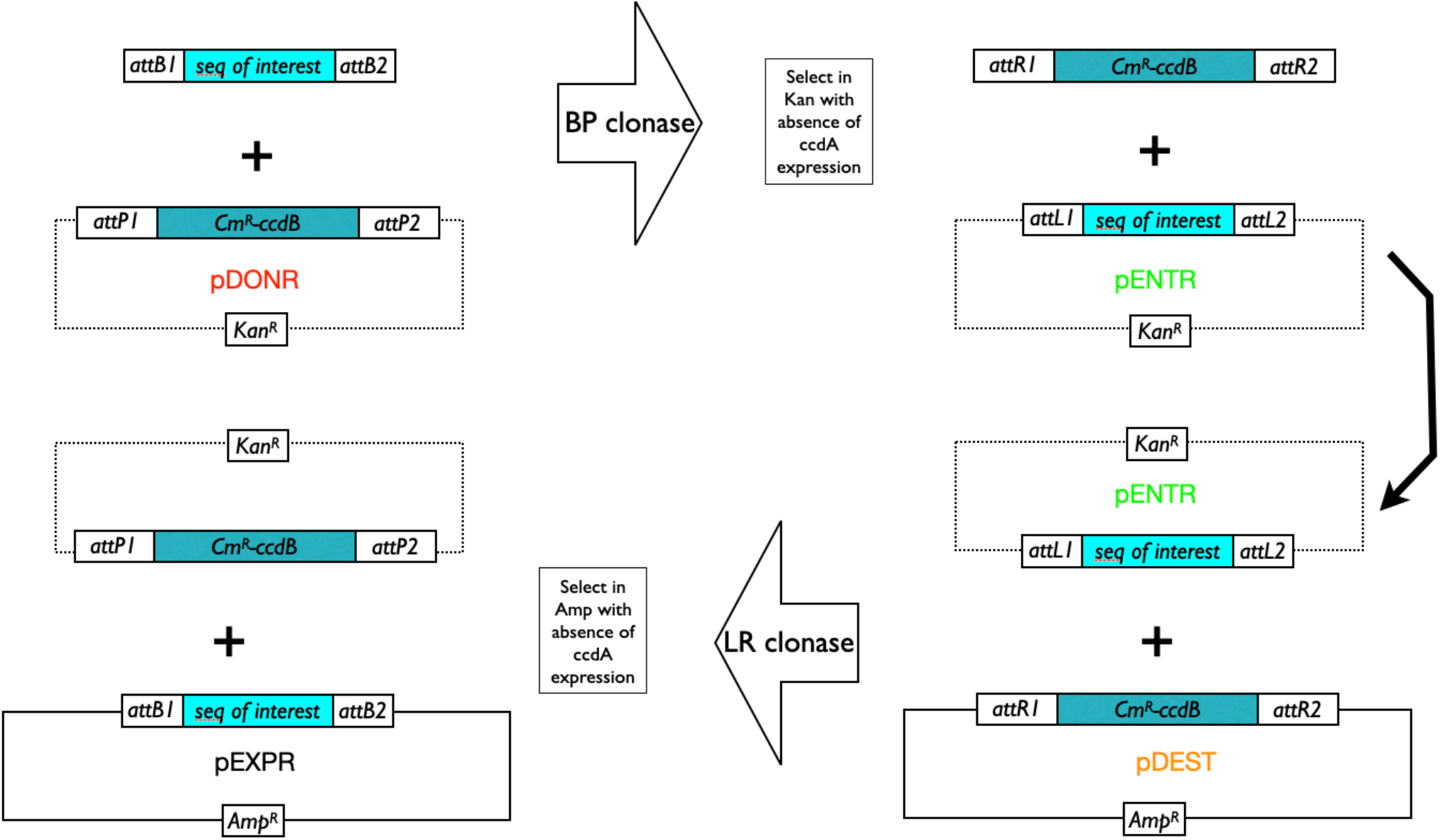
Principles of BP and LR Gateway cloning. Recombination specificity is dictated by the nature of the 7-base-pair core sequence, O, within each att site. A handful of att sequences with unique O-sequences have been selected to maximize recombination efficiency and minimize cross-reactivity between non-identical att sites. As such, a predefined order, orientation, and reading frame can be achieved by leveraging mutated lambda att-site variants with high specificity and virtually no crosstalk (orthogonality). (Upper panel) For example, attB1 recombines with attP1, but not with attP2 (and neither attP3, attP4, attP5, nor attP6). (Lower panel) Schematic of a typical Gateway LR cloning reaction, which combines a pENTR plasmid encoding a promoter and /or a sequence/gene of interest (GoI) with a pDEST plasmid harboring the rAAV replication (ITRs) and expression (WPRE, pA, etc.) motifs to be incorporated into the final pEXPR construct, ready for viral packaging. BP and LR clonase are commercially available mixtures of IHF and Int or IHF, Int, and Xis enzymes, respectively.

Optical interrogation of neural circuits places particularly stringent requirements on transgene expression. Depending on the application, experiments may require cell-type specificity, sparse or dense labelling, subcellular targeting of indicators or actuators, co-expression of multiple components, or precise tuning of expression density — and often simultaneously. Added to this is the rapidly expanding diversity of available indicators and actuators for any given modality, as illustrated by the proliferation of GEVI variants alone (ASAP 1–6, JEDI 1–3, FORCE1s, 1f, VADER1, Voltron1, 2, and others). For rAAV-mediated expression, Gateway cloning offers a practical solution: promoter elements, transgene sequences, and conditional expression cassettes can each be maintained as permanent entry clone libraries and recombined in any combination without repetitive subcloning.

These requirements become especially acute in the context of advanced optical acquisition methods that impose their own constraints on expression. Here, we illustrate this principle using ULoVE (Ultra-fast Local Volume Excitation), a two-photon microscopy method based on acousto-optic deflectors (AODs) that provides kHz-rate sampling with high signal-to-noise ratio in vivo and in acute slice preparations (Villette et al., 2019). Because ULoVE uses spatially extended or shaped excitation volumes, recordings are sensitive to out-of-focus fluorescence from neighboring structures, making precise control of expression sparsity, subcellular localization, and intensity essential prerequisites for high-quality data. As we show below, the Gateway approach is ideally suited to meet these requirements across a range of experimental contexts.

This article provides a comprehensive account of Gateway cloning methodologies specifically tailored for rAAV-mediated optical interrogation of neural circuits, addressing vector design principles, construction workflows, and quality control measures. As a specific demonstration of the approach, we present in vivo recordings from Purkinje cell dendrites in awake, behaving mice and optogenetic photostimulation of molecular layer interneurons in acute cerebellar slices — preparations that collectively demand cell-type specificity, sparse labelling, sub-millisecond temporal resolution, and precise density control — as these requirements illustrate the versatility and practical value of the Gateway pipeline when deployed alongside ULoVE.

## Results

### A combinatorial Gateway cloning library for optical neuroscience

To meet the expression demands of optical neuroscience applications, we constructed a modular Gateway cloning library comprising promoter entry clones, transgene entry clones, and AAV destination vectors designed for MultiSite LR recombination **(see Materials and Methods for details; Figure 3)**. Any promoter entry clone can be combined with any transgene entry clone in a single overnight reaction to generate a functional rAAV expression plasmid, without re-cloning either component. To date, in establishing this system, we have performed 105 viral productions spanning 9 promoter elements and more than 20 transgene configurations across multiple AAV capsids **(Figure 3, right panel; see Supplement to Figure 3 and Supplemental Table 1 for the full indexed production record of all 69 Gateway constructs to date, including capsid types and plasmid identifiers)**. This production record illustrates the practical combinatorial leverage of the Gateway approach: constructs that would each require independent cloning campaigns by conventional methods were generated here by simple recombination of existing library elements. The practical value of this modularity is demonstrated directly in the ULoVE recording experiments described below, where GCaMP6f, jRGECO1a, and JEDI2P-Kv were each conditionally expressed under matched, cell-type-specific conditions using a common Gateway-constructed L7::Cre driver virus **(see section ‘Optimized recordings of functional signals using ULoVE’)**.

**Figure 3.**
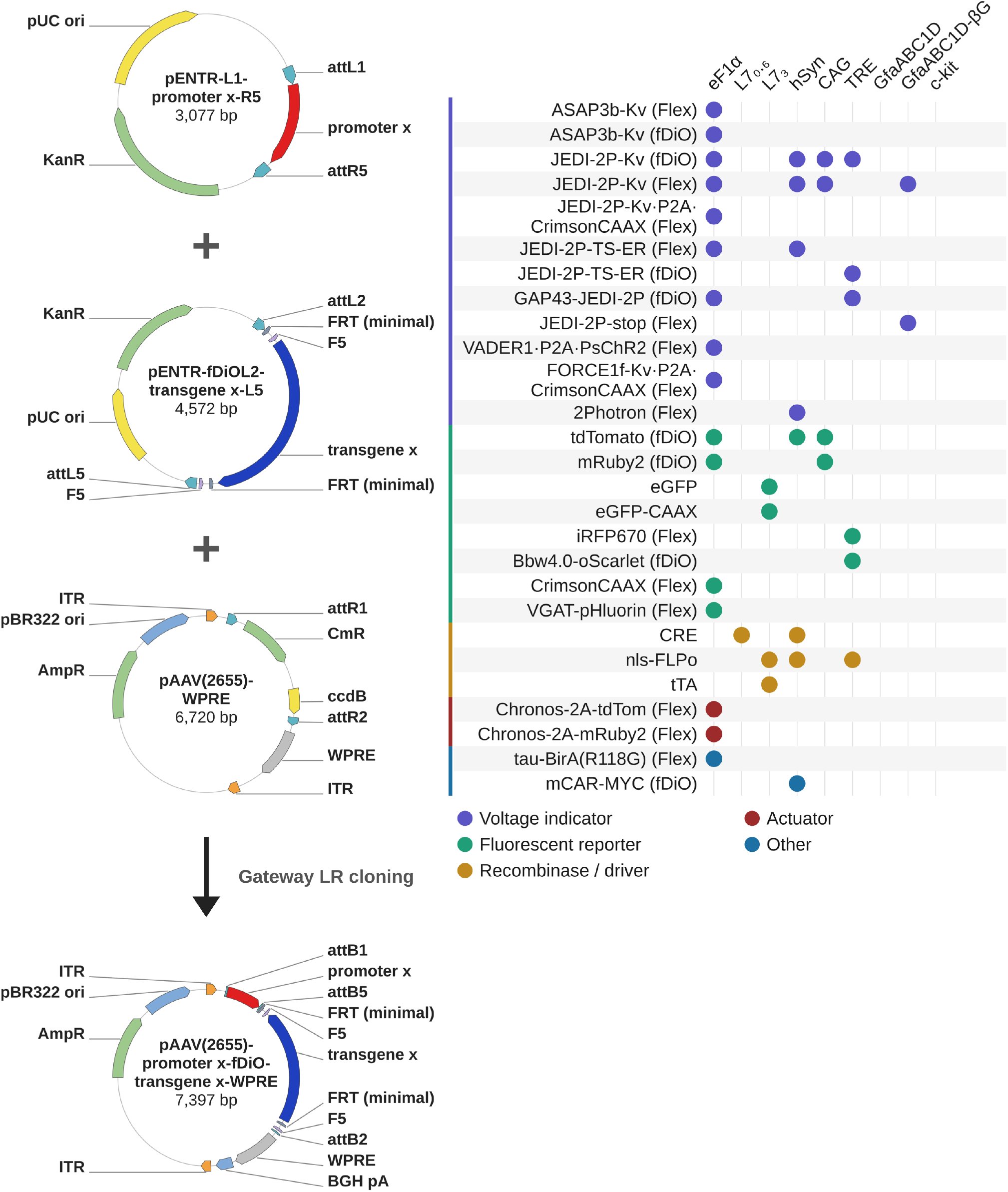
MultiSite Gateway cloning strategy for the generation of conditional rAAV expression constructs, and a combinatorial library of in-house pEXPR constructs that have been produced. **(Left panel)** Schematic of the three-component MultiSite LR cloning reaction used to produce all rAAV pEXPR plasmids in this study. Top to bottom: The promoter entry clone (pENTR-L1-promoter x-R5) carries the promoter element of interest flanked by attL1 and attR5 recombination sites. The transgene entry clone (pENTR-fDiOL2-transgene x-L5) carries the transgene of interest in an inverted, FLP-dependent (fDiO) configuration flanked by attL5 and attL2 recombination sites. The AAV destination vector (pAAV(2655)WPRE) contains AAV2 inverted terminal repeats (ITRs, orange) flanking attR1 and attR2 recombination sites, along with WPRE and BGH polyadenylation signal for efficient transgene expression, and a low-copy-number pBR origin of replication to minimize ITR instability during bacterial propagation. Gateway LR Clonase Plus enzyme mediates simultaneous recombination of all three components: the attL1/attR5 and attL5/attL2 site pairs insert the promoter and transgene at their defined positions and orientation within the destination attR1-attR2 vector, replacing the ccdB-CmR negative selection cassette. The resulting pEXPR construct (pAAV(2655)-promoter x-fDiO-transgene x-WPRE) contains the assembled expression cassette flanked by ITRs, ready for rAAV production. In the example shown, the transgene remains in the inverted (non-coding) orientation until FLP recombinase activity catalyzes permanent reversion to the sense orientation. Analogous reactions using Flex (CRE-dependent) transgene entry clones, unconditional transgene entry clones, or alternative destination vector backbones generate the full range of constructs described in this study. **(Right panel)** Dot matrix: Representative examples summarizing the combinatorial Gateway cloning library of in-house pEXPR constructs produced. Rows indicate transgene entry clone identities, grouped by functional category: voltage indicators (purple), fluorescent reporters (green), recombinase/driver constructs (gold), actuators (red), and other constructs (blue). Columns indicate promoter entry clone identities: eF1α, L7_0·6_, L7_3_, hSyn, CAG, TRE, GfaABC1D, GfaABC1D-βG, and c-kit. Each filled dot indicates a promoter–transgene combination for which at least one rAAV production was completed. The full indexed production record of all 69 Gateway constructs to date, including capsid types and plasmid identifiers, is provided in **Supplement to Figure 3** and **Supplemental Table 1**)

### Application to sparse and cell-type-specific expression of reporters

#### Capsid type considerations

Typically, inverted terminal repeats (ITRs) from AAV type 2 are included in the final destination/expression vectors and are compatible with viruses being pseudotyped with virtually all naturally occurring and engineered capsids, each exhibiting somewhat distinct tropism profiles and transduction capacities (Aschauer et al., 2013; Jang et al., 2023). For most naturally occurring capsids, tropism is governed by the cellular entry receptor, including heparan sulfate proteoglycan (types 2, 3, and 6), α2,3-sialylated glycoproteins (types 1, 4), O-linked glycoproteins (type 5), N-linked glycoproteins (type 6), laminin receptors (type 8), and galactose (type 9). To date, more than 10 natural capsids have been used for rAAV production, plus engineered variants including DJ and DJ/8, a shuffled capsid type engineered from AAVs 2, 8, and 9 to give broader tropism (Grimm et al., 2008). Types rh10 and rh32.33 have been isolated from rhesus macaques and escape neutralizing antibodies. Specialized properties also exist: AAV1 has been shown to support effective transsynaptic anterograde transport for targeted conditional expression (Zingg et al., 2017, 2020; Beltramo and Scanziani, 2019); AAV5 exhibits diffuse spread and pronounced astrocyte tropism, beneficial for large-volume transduction (Davidson et al., 2000); rAAV2-retro enables retrograde access to projection neurons (Tervo et al., 2016); and AAV-PHP.eB demonstrates enhanced brain transduction following intravenous or intraventricular administration, although BALB/cJ mice are resistant due to polymorphisms in its receptor Ly6a (Huang et al., 2019). AAV-PHP.S supports peripheral nervous system transduction (Chan et al., 2017).

#### Cell-type specific and/or sparse expression strategies

Gateway destination vectors can incorporate multiple strategies for cell-type-specific expression. CRE or FLPo recombinase-dependent vectors utilize inverted coding sequences flanked by heterotypic pairs of lox or frt recombination sites that undergo recombination only in recombinase-expressing cells (Atasoy et al., 2008; Sohal et al., 2009; Fenno et al., 2014). In combination with mouse lines genetically programmed to drive recombinase expression in defined cell types — for example, interneurons expressing parvalbumin (PV; Hippenmeyer et al., 2005(Hippenmeyer et al., 2005)), somatostatin (SST), vasoactive intestinal peptide (VIP), or cholecystokinin (CCK), among others (Taniguchi et al., 2011(Taniguchi et al., 2011)), or GABAergic neurons expressing glutamate decarboxylase 67 (GAD67; Fuchs et al., 2001; Tolu et al., 2010(Fuchs et al., 2001; Tolu et al., 2010)) — the GoI coding sequence is reverted to the functional orientation only in recombinase-expressing cells. This intersectional strategy can also be leveraged to achieve sparse GoI expression by delivering the recombinase via a second AAV co-injected at a limiting titer relative to the conditional indicator virus. Using the recombinase-driver virus carrying a pan-neuronal (hSyn) or cell-type-specific (CaMKII) promoter and injected at titers 50–100-fold lower than the GoI expressing virus has proven invaluable for imaging-based studies where dense expression would preclude high signal-to-noise recording from GECIs or GEVIs (Sheffield and Dombeck, 2015; Chan et al., 2017; Villette et al., 2019). As demonstrated directly in the ULoVE-based Purkinje cell recordings described below, lowering the titer of the CRE-driver AAV over a 100-fold range (∼1×10^10-12^ GC/ml, co-injected against ∼1×10^11-12^ GC/ml indicator virus) produced a graded reduction in labelling density sufficient to resolve individual dendritic arbors — a prerequisite for high signal-to-noise ULoVE recordings in which neuropil contamination from neighboring structures directly limits optical recording fidelity **(Figure 6B)**.

The Gateway approach also enables sparse and cell-type-specific expression by leveraging the myriad recombinase driver lines of mice available as BAC transgenic or knock-in strains, in combination with sophisticated intersectional viral strategies. **Figure 4** illustrates this principle using a triple-intersectional approach in a vGlut2::CRE; GlyT2::eGFP double-transgenic mouse, in which the GlyT2::eGFP allele provides a constitutive reporter of glycinergic neurons — including inhibitory deep cerebellar nuclei (DCN) neurons and cerebellar cortical Golgi and Lugaro cells — independent of the viral strategy. CRE recombinase expressed in vGlut2+ excitatory DCN neurons drives expression of the two co-injected CRE-dependent viruses. The first, AAV1.eF1α.DiO.nls-FLPo (virus ENS 008, produced in-house from Addgene 87306), provides nuclear-localized FLPo recombinase. The second, AAV1.ihSyn1.DiO.tTA (Addgene 99121-AAV1) provides the tetracycline transactivator tTA. A third, Gateway-cloned virus, AAV1.TRE.fDiO.tdTomato (virus ENS 068, pEXPR A-183) was assembled by MultiSite LR recombination of a TRE promoter entry clone with a fDiO-tdTomato transgene entry clone. Expression of tdTomato from this construct requires both tTA activity to drive transcription from the TRE promoter and FLPo activity to invert the fDiO cassette into the sense orientation. This dual requirement, combined with the limiting titer of the CRE-dependent FLPo-expressing virus, restricts labelling to a sparse subset of vGlut2+ neurons, while TRE-driven amplification ensures strong fluorescence suitable for axonal projection tracing. Using this strategy, we traced vGlut2+ DCN axonal projections into the cerebellar cortex, where terminations were concentrated preferentially at the boundary between the granule cell layer and the molecular layer, consistent with previous descriptions of DCN→cortex nucleocortical projections (Houck and Person, 2015; Gao et al., 2016). Volumetric analysis of CUBIC-cleared tissue confirmed this laminar preference quantitatively, with mean axonal signal intensity 40% higher in the combined Purkinje cell/superficial granular layer than across the lobe as a whole (**Supplement to Figure 4**).

**Figure 4.**
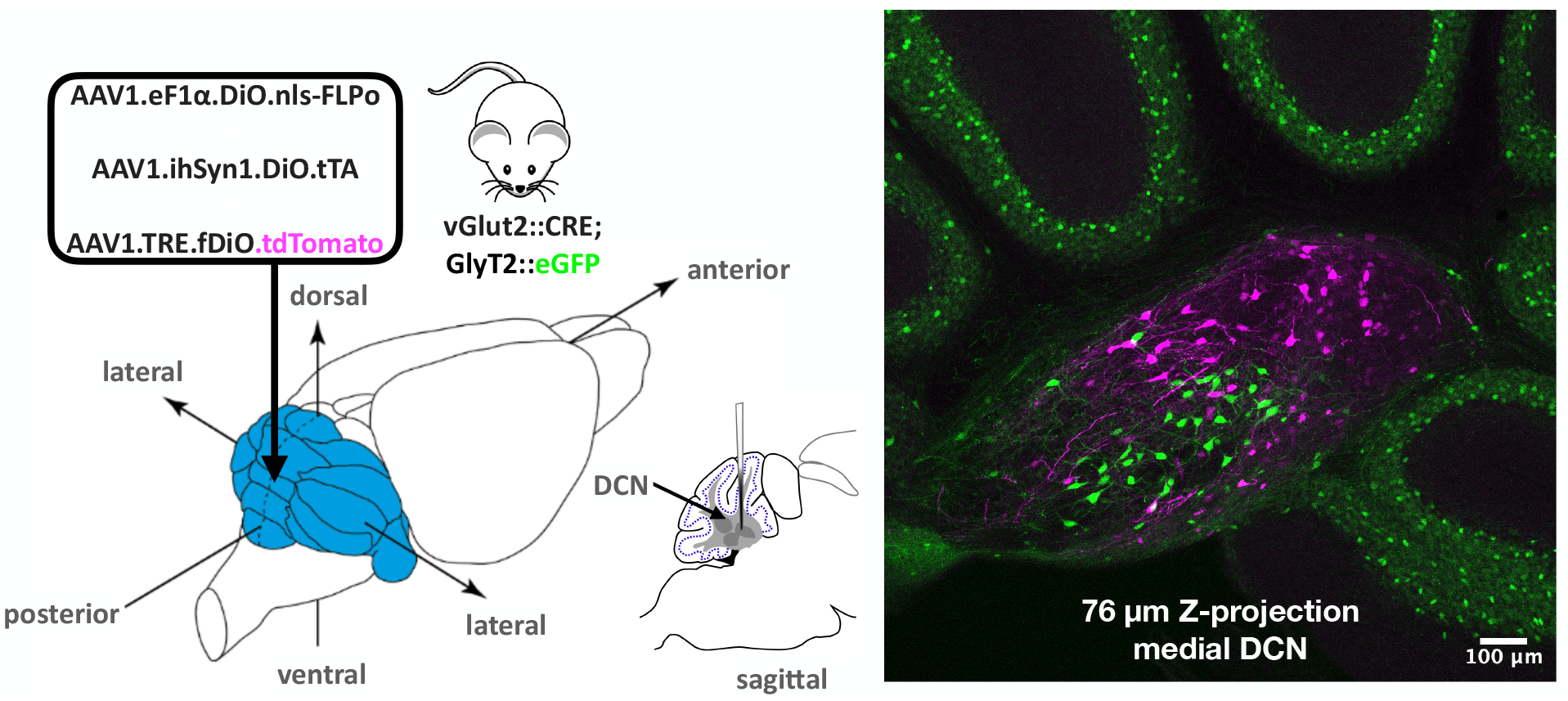
Sparse, strong, and cell-type-specific expression in the medial deep cerebellar nuclei using a triple-intersectional Gateway-cloned viral strategy. **(Left)** Experimental design. A mixture of three AAVs (boxed) was injected into the cerebellum (blue) of a vGlut2::CRE; GlyT2::eGFP double-transgenic mouse, with the pipette targeted to the medial DCN (inset, sagittal view; medial, interposed and lateral DCN represented in dark grey; stereotaxic coordinates in Methods). CRE recombinase expressed in vGlut2+ excitatory DCN neurons drives expression of the two co-injected CRE-dependent viruses AAV1.eF1α.DiO.nls-FLPo and AAV1.ihSyn1.DiO.tTA. A third Gateway-cloned virus, AAV1.TRE.fDiO.tdTomato (virus ENS 068, pEXPR A-183), assembled by MultiSite LR recombination of a TRE promoter entry clone with a fDiO-tdTomato transgene entry clone, requires both tTA-driven TRE activation and FLPo-mediated inversion of the fDiO cassette for tdTomato expression. This restricts labelling to vGlut2+ neurons in which all three conditions are satisfied. A limiting titer of the CRE-dependent FLPo-expressing virus further restricts the labelled population to a sparse subset. **(Right)** Representative fluorescence image (76 µm z-projection) showing sparse but strong tdTomato expression (magenta) in vGlut2+ excitatory neurons of the medial DCN, with individual axonal projections clearly resolvable for anatomical tracing (see Supplement to Figure 4), alongside GlyT2::eGFP-labelled inhibitory DCN neurons (green). GlyT2::eGFP also labels glycinergic Golgi and Lugaro cells of the cerebellar cortex, providing anatomical context within and around the injected structure. Scale bar: 100 µm.

Achieving more precisely sculpted expression by using AAVs to drive recombinase expression from promoters or enhancer elements active in transcriptionally defined neuronal subtypes has proven challenging but is feasible in certain cases (Dimidschstein et al., 2016; Graybuck et al., 2021). Two examples we have found successful using the Gateway pipeline **(Figure 5)** are previously characterized fragments, 0.6kb (L7_0.6_), 1kb (L71), and 3kb (L7_3_) of the L7/pcp2 promoter for Purkinje cell-specific expression (Oberdick et al., 1990; El-Shamayleh et al., 2017; Nitta et al., 2017) and a fragment of the c-kit oncogene promoter for molecular layer interneuron expression with limited Golgi cell co-labelling (Cairns et al., 2003; Schilling and Oberdick, 2009; Amat et al., 2017; Villette et al., 2019). Notably, we have found that these expression strategies function equally well in rats (Jalil and Bradley, unpublished; and (Figueroa et al., 2023)).

**Figure 5.**
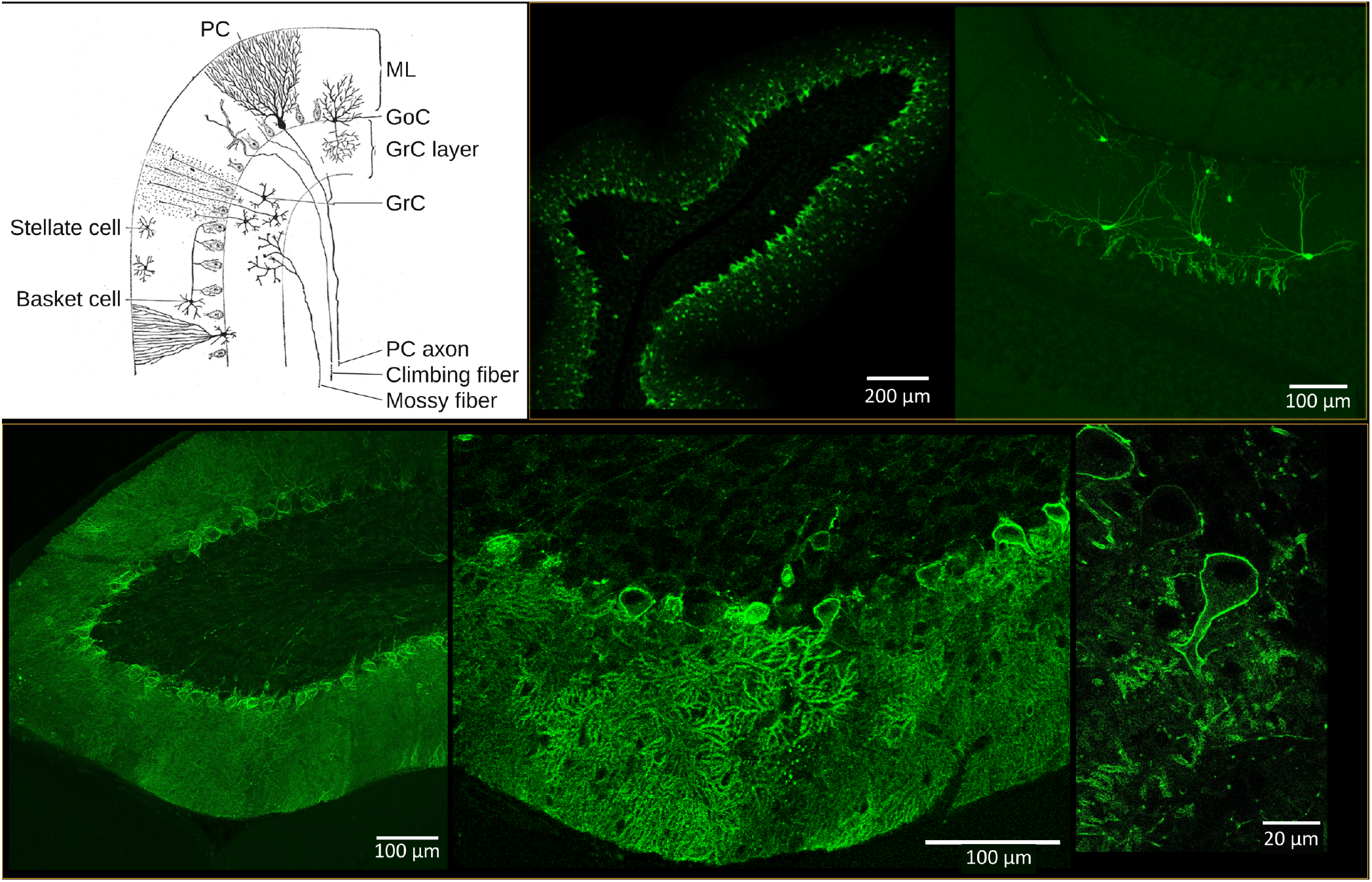
Cell-type-specific rAAV CRE and FLPo drivers across the folial architecture of the cerebellar cortex. **(Upper panel, left)** Schematic of the cerebellar cortex after Ramón y Cajal, showing Purkinje cells (PC), the molecular layer interneurons (MLIs) basket and stellate cells, Golgi cells (GoC) and granule cells (GrC), the molecular layer (ML) and the granule cell layer (GrC layer), and the PC axon, climbing fiber and mossy fiber. **(Upper panel, middle and right)** CRE-conditional expression of GCaMP6f in MLIs using the c-kit driver injected at different titers to obtain denser **(middle: kit::CRE injected at 7.7×10**^**12**^ **GC/ml)** or sparser **(right: kit::CRE injected at 7.7×10**^**10**^ **GC/ml)** expression. **(Lower panel**) FLPo-conditional expression of ASAP3b-Kv in Purkinje cells using AAV1.L7_3_.nls-FLPo as driver. Panels show successive magnifications of the same field. Full injection and viral parameters are described in Methods (“Stereotaxic viral injections for anterograde tracing of DCN projections (Figure 4) and validation of cell-type-specific rAAV drivers (Figure 5))”. Note that ASAP3b expression is also in Purkinje cell dendrites despite having the Kv2.1 PRC targeting motif.

### Complex vector assembly by MultiSite Gateway cloning

Many optical neuroscience applications require expression constructs that combine multiple functional elements — a cell-type-specific promoter, a conditional transgene cassette, and in some cases a second reporter or actuator — in a single viral vector. Our MultiSite Gateway system assembles such constructs in a single LR reaction by exploiting the orthogonality of attL1/attR5 and attL5/attL2 recombination site pairs, which ensures that the promoter and transgene entry clones are inserted sequentially at defined positions within the destination vector without cross-reactivity (Sasaki et al., 2004); **Figure 3**). This approach has proven particularly powerful for the rapid generation of recombinase-conditional vectors, in which the transgene entry clone carries an inverted coding sequence flanked by heterotypic lox or frt site pairs; a single LR reaction converts this entry clone into a CRE- or FLPo-dependent expression construct ready for AAV packaging, reducing construction time from weeks to days. As illustrated in **Figure 4**, this framework also supports inducible conditional expression: a TRE promoter entry clone combined with a fDiO-tdTomato transgene entry clone in a single MultiSite LR reaction produced pEXPR A-183 (virus ENS 068), the reporter component of a triple-intersectional strategy for sparse, strong axonal projection tracing of excitatory DCN neurons (see section “Application to sparse and cell-type-specific expression of reporters”). The same framework accommodates bicistronic constructs in which two coding sequences are separated by a P2A ribosomal-skipping peptide, enabling co-expression of, for example, an optogenetic actuator and an anatomical marker from a single vector. Technical considerations for MultiSite reactions — including stoichiometric ratios, extended incubation times, and screening strategies — are detailed in the Materials and Methods and in the Discussion.

### Optimized recordings of functional signals using ULoVE

To demonstrate how the versatility of the Gateway cloning pipeline serves the characterization of neuronal dynamics across multiple timescales, we examined Purkinje cell dendritic activity in awake, head-fixed behaving mice using three genetically encoded fluorescent indicators. Purkinje cell dendrites receive powerful excitatory input from climbing fibers that evoke large, stereotyped dendritic depolarizations, making them an ideal testbed for comparing calcium and voltage indicator performance at the temporal limits of in vivo optical recording. Previous studies of Purkinje dendritic dynamics have relied on dense labelling strategies and, in most cases, anaesthetized animals (Ozden et al., 2008; Mukamel et al., 2009; Roome and Kuhn, 2018). Here, we combined Gateway-enabled cell-type-specific and sparse expression with ULoVE two-photon excitation to achieve optical recordings at a sampling rate of ∼5 kHz from sparsely labelled dendrites in awake mice, permitting resolution of near-onset voltage and calcium dynamics with sub-millisecond precision.

#### Gateway-enabled expression strategies for Purkinje cell indicators

Selective and sparse expression of optical indicators in Purkinje cells was achieved using several complementary viral strategies built on Gateway-cloned driver viruses **(Figure 6A**). For calcium imaging, GCaMP6f and jRGECO1a were each expressed using a CRE-dependent approach: AAV1.L7_0·6_.CRE was co-injected at limiting dilution with either AAV1.CAG.Flex.GCaMP6f or AAV1.CAG.Flex.NES-jRGECO1a, respectively (strategies ➀ and ➁). For voltage imaging, two independent strategies were employed. Strategy ➂ used AAV1.L7_3_.CRE co-injected with AAV1.eF1α.Flex.JEDI2P-Kv (Liu et al., 2022) for single-recombinase conditional expression, while an alternative dual-intersectional approach combined AAV1.L7_3_.nls-FLPo and AAV1.L7_3_.tTA with AAV1.TRE.fDiO.JEDI2P-Kv to enhance indicator expression through TRE-driven amplification while preserving sparse labelling through FLP-dependent conditionality. Each strategy was delivered to a separate cohort of mice (GCaMP6f, n = 2; jRGECO1a, n = 1; JEDI2P-Kv, n = 2), so no cell co-expressed more than one indicator, and calcium and voltage were not recorded simultaneously. The recordings in Figure 6D–J therefore come from independent dendrites, compared across populations of climbing fiber-evoked events acquired with the same ULoVE recording approach. The driver viruses (AAV1.L7_0·6_.CRE, AAV1.L7_3_.CRE, AAV1.L7_3_.nls-FLPo, AAV1.L7_3_.tTA) and the two JEDI2P-Kv vectors were generated using the MultiSite Gateway pipeline described above, enabling rapid systematic variation of promoter length (L7_0·6_ versus L7_3_), recombinase type (CRE, FLPo), transactivator (tTA), and JEDI2P-Kv expression configuration without re-cloning from scratch; GCaMP6f and jRGECO1a were obtained from Addgene (see Methods). Lowering the titer of the CRE-driver AAV over a 100-fold range (1×10^12^ to 1×10^10^ GC/ml co-injected against a fixed 1×10^11^ GC/ml indicator virus) produced a graded reduction in labelling density, such that individual dendritic arbors could be resolved even at considerable distance from the injection site (**Figure 6B)**. This precise density control — essential for ULoVE recordings where neuropil background directly limits signal-to-noise — is a direct consequence of the Gateway pipeline’s capacity to generate matched, interchangeable recombinase-driver and reporter constructs efficiently.

**Figure 6.**
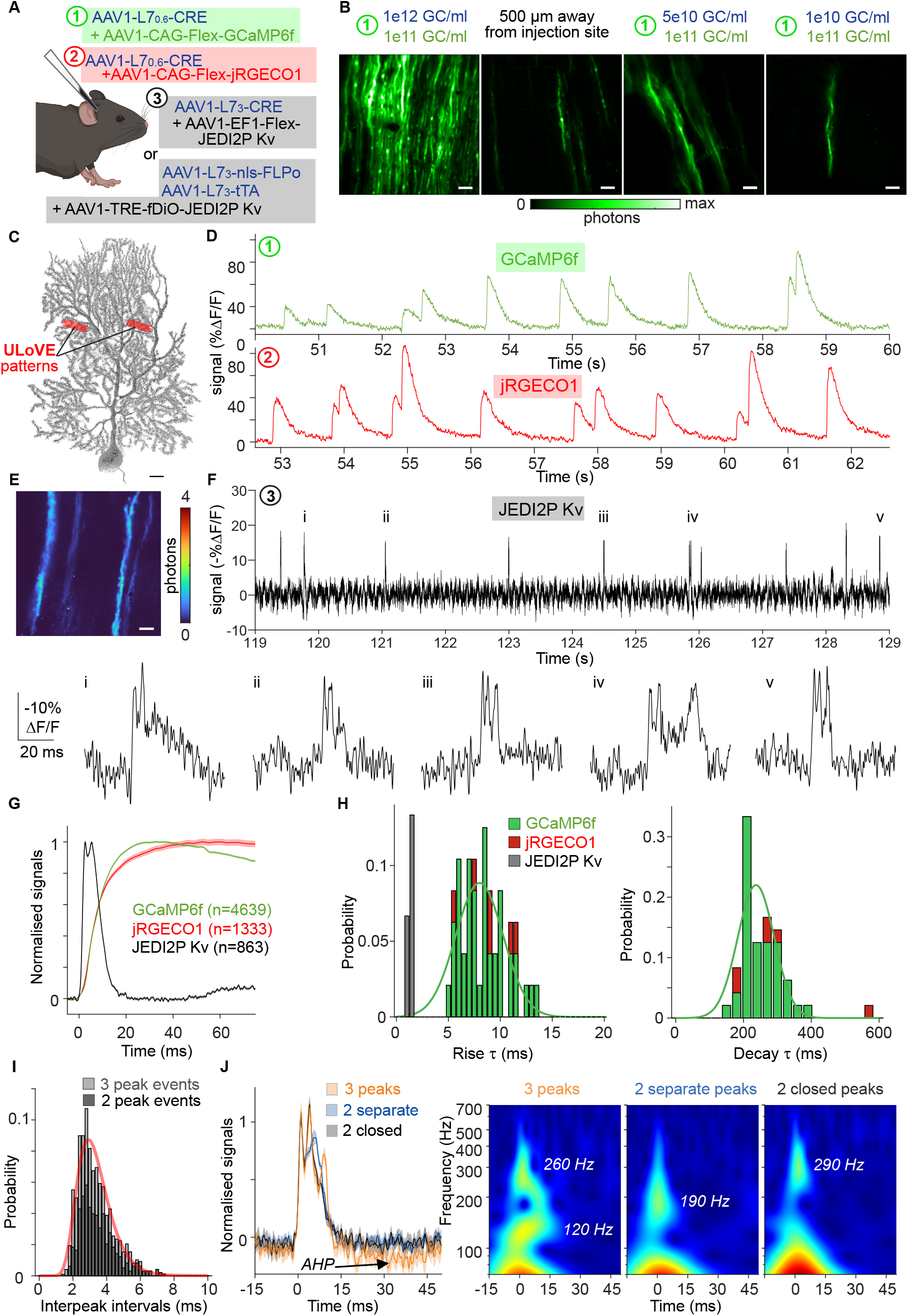
In vivo recording of Purkinje cell dendritic activity with genetically encoded calcium and voltage indicators using ULoVE two-photon excitation. **(A)** Schematic of the viral injection strategies used to selectively express fluorescent indicators in Purkinje cells using Gateway-cloned driver viruses; the GCaMP6f and jRGECO1a constructs in ➀ and ➁ were obtained from Addgene. Each strategy was used in separate animals. Numbered circles (➀–➂) indicate the color-coded strategies referenced throughout: ➀ GCaMP6f (green), ➁ jRGECO1a (red), ➂ JEDI2P-Kv (black). For JEDI2P-Kv, two strategies were designed — a single-recombinase CRE/lox approach analogous to the calcium indicator strategies, and a dual-intersectional approach combining TRE-driven expression amplification with FLP-dependent conditionality to enhance sparse labelling. **(B)** Representative two-photon fluorescence images of Purkinje cell dendrites labelled using strategy ➀ at three CRE-driver dilutions (1×10^12^, 5×10^10^, and 1×10^10^ GC/ml co-injected against a fixed 1×10^11^ GC/ml indicator virus). Reducing the CRE-driver titer progressively decreases labelling density; sparse individual arbors can be resolved 500 µm from the injection site. Images displayed on a photon-count scale (0 to max; max = 12 except for the 1×10^10^ GC/ml condition, where max = 5). Scale bars: 20 µm. **(C)** Three-dimensional reconstruction of a Purkinje cell (grayscale) with two representative ULoVE excitation volumes (red) positioned on the dendritic arbor. Scale bar: 10 µm. **(D)** Representative in vivo calcium transients from Purkinje cell dendrites expressing GCaMP6f (strategy ➀, top, green) and jRGECO1a (strategy ➁, bottom, red), showing normalized ΔF/F over time. Repeated stereotyped events correspond to climbing fiber-evoked dendritic calcium transients. Traces smoothed with a 3 ms Gaussian kernel for display. **(E)** Representative two-photon fluorescence image of Purkinje cell dendrites expressing JEDI2P-Kv (strategy ➂). Scale bar: 20 µm. **(F)** Voltage trace recorded from a JEDI2P-Kv-expressing Purkinje cell dendrite showing spontaneous dendritic events (labelled i–v), displayed as –ΔF/F so that membrane depolarizations appear as positive deflections. Trace smoothed with a 0.5 ms Gaussian kernel. Bottom: expanded views of individual events i–v from the raw trace, illustrating the diversity of event morphology, including two- and three-peak spikelets. **(G)** Average normalized waveforms for GCaMP6f (n = 4639 events, 42 dendrites), jRGECO1a (n = 1333 events, 6 dendrites), and JEDI2P-Kv (n = 863 events, 6 dendrites), aligned to event onset. Shaded areas: ±SEM. Note the markedly faster kinetics of the voltage indicator. **(H)** Distributions of exponential rise time constants (τ_rise_, left) and decay time constants (τ_decay_, right) estimated per dendrite from average waveforms of temporally isolated onset-aligned events. Solid green curves: Gaussian fits to summed distributions. **(I)** Distribution of inter-peak intervals for JEDI2P-Kv dendritic events classified as 2-peaks (dark grey, n = 521) or 3-peaks (light grey, n = 229). Log-normal fit (red curve) reveals a skewed distribution with excess around 2.5 and 6 ms. **(J)** Left: average normalized waveforms for 3-peaks (orange, n = 16), 2-separate-peak (blue, n = 27), and 2-closed-peaks (grey, n = 19) event categories, selected for temporal homogeneity. Shaded areas: ±SEM. An afterhyperpolarization (AHP, arrow) is present in 3-peak events and absent from 2-peak events. Right: time-frequency spectrograms (Morlet wavelet analysis) for each category, revealing oscillatory components at 290 Hz (2-closed-peaks), 190 Hz (2-separate-peaks), and 260/120 Hz (3-peaks).

#### ULoVE recording of calcium and voltage transients

ULoVE excitation patterns were positioned in the middle of the dendritic arbor **(Figure 6C)**, targeting the region most likely to capture climbing fiber-evoked events. Both GCaMP6f and jRGECO1a reliably reported repeated stereotyped calcium transients consistent with spontaneous climbing fiber activation **(Figure 6D)**, with high signal-to-noise at ∼5 kHz sampling. Voltage recordings with JEDI2P-Kv revealed rapid dendritic depolarizations with a rich waveform structure **(Figure 6F)**. The raw voltage trace shows discrete transient events of varying duration and morphology (events i–v, **Figure 6F**), with expanded single-event traces revealing sub-millisecond structure that would be entirely unresolvable at conventional calcium imaging frame rates.

#### Comparative kinetics of calcium and voltage signals

Averaged onset-aligned transients from large event populations (GCaMP6f: n = 4639 events, 42 dendrites; jRGECO1a: n = 1333 events, 6 dendrites; JEDI2P-Kv: n = 863 events, 6 dendrites) illustrated the markedly different timescales of calcium versus voltage signals **(Figure 6G)**. Exponential time constants for rise and decay were estimated per dendrite from temporally isolated events **(Figure 6H)**. Strikingly, both calcium indicators exhibited rise time constants (τ_rise_) that were substantially faster than those previously reported in cortical neurons (Chen et al., 2013; Dana et al., 2016; Park et al., 2021), with a mean τ_rise_ of 8.44 ± 2.17 ms. This acceleration likely reflects the exceptionally large and rapid calcium influx associated with climbing fiber-evoked dendritic depolarizations, which may drive calcium indicators closer to saturation on a faster timescale than the sparser calcium entry during cortical spiking, and is consistent with earlier work in cerebellar slices showing that climbing fiber-induced calcium dynamics occur within 10 ms and can drive multiple dendritic calcium spikes (Miyakawa et al., 1992; Otsu et al., 2014). Notably, GCaMP6f and jRGECO1a showed comparable rise kinetics under these conditions, suggesting that in this regime, the underlying biological calcium dynamics, rather than indicator binding rates, are the primary determinant of response speed. Despite this acceleration relative to cortical benchmarks, calcium rise remained approximately 6.7-fold slower than the voltage depolarization rise measured with JEDI2P-Kv (τ_rise_ = 1.25 ± 0.24 ms), underscoring the fundamental kinetic advantage of direct voltage sensing. Calcium decay was substantially prolonged (τ_decay_ = 254 ± 71.9 ms), consistent with the slow unbinding kinetics of GECIs and sustained calcium extrusion dynamics following climbing fiber activation.

#### Dendritic spikelets resolved by ULoVE voltage imaging in awake mice

Examination of individual JEDI2P-Kv events at full ∼5 kHz temporal resolution revealed a complex waveform structure within each climbing fiber-evoked depolarization: a subset of events exhibited two or three distinct sub-peaks, or “spikelets”, superimposed on the rising phase of the voltage transient **(Figure 6F, insets i–v)**. Such spikelets have been examined using somatic intracellular recording (Maruta et al., 2007), dendritic patch clamp in anesthetized preparations (Kitamura and Häusser, 2011), and the synthetic voltage dye ANNINE-6+ in awake mice (Roome and Kuhn, 2018), but have not, to our knowledge, been resolved by optical voltage recording with GEVIs in awake behaving animals. Analysis of inter-peak intervals across 2-peak (n = 521) and 3-peak (n = 229) events revealed an asymmetric distribution with a broad primary peak around 2–4 ms and a secondary excess of events around 6 ms relative to the log-normal fit (Figure 6I), suggesting that spikelets can occur at discrete inter-event intervals, but with some jitter, reflecting the underlying oscillatory or resonant properties of climbing fiber–Purkinje cell interactions. Selecting temporally homogeneous event subsets to characterize average waveforms and dominant oscillatory content (**Figure 6J**) revealed that 2-peak events with closely spaced peaks showed a dominant frequency of ∼290 Hz, while those with more separated peaks showed ∼190 Hz. Three-peak events displayed frequency adaptation, decelerating from ∼260 Hz to ∼120 Hz across successive peaks, and were additionally distinguished by a clear afterhyperpolarization (AHP) following the final peak — a feature absent from all 2-peak event categories (**Figure 6J, left**). Time-frequency spectrograms confirmed these discrete oscillatory components across event categories **(Figure 6J, right)**. The selective presence of an AHP in 3-peak events may reflect the recruitment of calcium-activated potassium conductances (Womack et al., 2004): the larger calcium influx associated with a more prolonged climbing fiber depolarization may activate SK or BK channels sufficiently to produce a measurable hyperpolarization. The varying inter-peak intervals and discrete frequency content across event categories may further suggest that successive dendritic spikelets arise from distinct resonance modes within the olivocerebellar circuit or from active dendritic conductances that shape climbing fiber signal propagation along the Purkinje cell arbor (Otsu et al., 2014).

Taken together, these results demonstrate how the Gateway cloning pipeline — by enabling rapid modular construction of cell-type-specific, density-tunable expression constructs across multiple recombinase systems — provided the expression conditions necessary for ULoVE to resolve sub-millisecond dendritic voltage dynamics alongside millisecond-scale calcium kinetics in awake behaving mice. The capacity to compare multiple indicator classes in matched experimental conditions, made possible by the combinatorial efficiency of Gateway cloning of driver and reporter constructs, is a key strength of this integrated approach.

### Optimized photoactivation of neurons using ULoVE

To illustrate a further application of the Gateway cloning pipeline, we examined whether ULoVE-based two-photon photostimulation could achieve cell-specific, sub-millisecond optogenetic activation of cerebellar molecular layer interneurons (MLIs) in acute brain slices. MLIs — of which basket cells represent the predominant subtype in the cerebellar molecular layer — provide powerful feedforward inhibition onto Purkinje cells and are therefore a key target for dissecting inhibitory microcircuit connectivity. Achieving single-cell optogenetic control of MLIs requires both cell-type-specific opsin expression and a photostimulation strategy with spatial resolution below the typical MLI soma diameter of 9–10 µm.

#### Gateway-enabled cell-type-specific expression of ChR2(H134R)-YFP in MLIs

Cell-type-specific expression of ChR2(H134R)-YFP in MLIs was achieved by co-injection of two Gateway-cloned viruses: AAV1.kit.CRE (virus ENS 001), which restricts CRE recombinase expression to MLIs via a 2.2 kb fragment of the c-kit promoter, and AAV1.DiO.hChR2(H134R)-YFP.WPRE.HGHpA (Addgene 20298), which drives conditional opsin expression only in CRE-expressing cells **(Figure 7A)**. As shown in **Figure 7B (bottom)** and previously in **Figure 5**, this strategy produces robust, cell-type-specific fluorescent labelling of MLIs throughout the cerebellar molecular layer. Acute cerebellar slices were prepared 15 days post-injection for combined whole-cell voltage-clamp recording and two-photon photostimulation.

**Figure 7.**
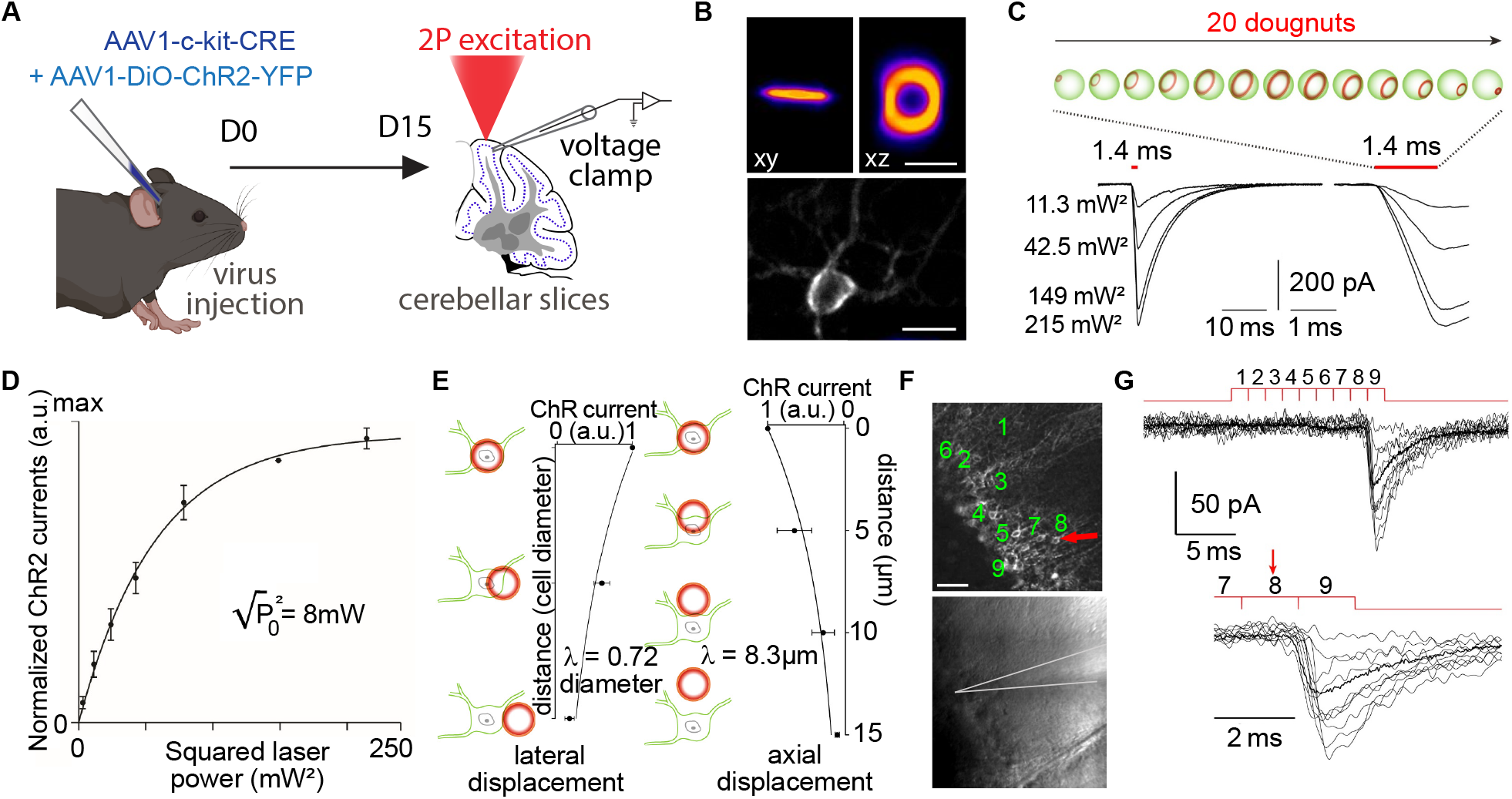
Two-photon multi-doughnut optogenetic photostimulation of ChR2(H134R)-YFP-expressing molecular layer interneurons in acute cerebellar slices. **(A)** Experimental timeline. Mice were injected with a CRE-dependent ChR2(H134R)-YFP construct together with the c-kit CRE driver (D0), and acute cerebellar slices were prepared at D15 for whole-cell voltage-clamp recordings combined with two-photon photostimulation. Full injection and viral parameters are described in Methods (“Cerebellar slice preparation and electrophysiology”). **(B)** Top: two-photon fluorescence images, shown rotated by 45° relative to the indicated plane (left, xy; right, xz), of the scanned doughnut excitation patterns. Scale bars: 10 µm. Bottom: representative two-photon fluorescence image of a ChR2(H134R)-YFP-expressing molecular layer interneuron (basket cell subtype). Scale bar: 10 µm. **(C)** Left: schematic of the multi-doughnut excitation pattern (20 doughnuts, 70 µs per doughnut, 1.4 ms total per cell). Right: representative ChR2(H134R) photocurrents recorded in whole-cell voltage clamp at increasing squared laser powers (11.3, 42.5, 149, and 215 mW^2^), with expanded timescale inset. Note that photocurrents were initiated before the full 20-doughnut sequence was completed, confirming sub-millisecond photoexcitation efficiency. **(D)** Input–output curve of normalized photocurrent amplitude as a function of squared laser power. Half-saturation power √P_0_^2^ = 8 mW. Data shown as mean ± SEM. **(E)** Spatial resolution of two-photon doughnut optogenetic activation. Photocurrent amplitude as a function of lateral (left) or axial (right) displacement of the ULoVE focus relative to the MLI soma. Lateral resolution constant: 0.72 cell diameters; axial resolution: 8.3 µm. **(F)** Top: two-photon fluorescence image of ChR2(H134R)-YFP-expressing MLIs; numbered cells (1–9) indicate the nine sequentially photostimulated cells. Red arrow indicates the presynaptically connected cell (MLI #8). Bottom: infrared Dodt contrast image showing the recorded Purkinje cell at the tip of the patch pipette. Scale bar: 50 µm. Photocurrents were probed at a revisit cycle frequency of 710 Hz, with each MLI receiving a 1.4 ms ULoVE excitation per cycle. **(G)** Evoked IPSCs recorded at the Purkinje cell soma from each of the nine sequentially photostimulated MLIs shown in (F). Only MLI #8 (red arrow; right inset) generated a reliable postsynaptic response, with a mean IPSC amplitude of 46 pA across 10 trials and one failure.

#### ULoVE doughnut-pattern photostimulation of single MLIs

The ULoVE excitation strategy used here differs from the extended-volume patterns used for the calcium and voltage recordings described above. Instead of using the acousto-optic deflectors to produce an elongated excitation volume matched to a dendritic arbor, individual MLI somata were photostimulated with doughnut-shaped excitation patterns generated by circularly scanning a diffraction-limited focus. (Villette et al., 2019). Given the small, approximately spherical geometry of MLI somata and the absence of motion artefacts in acute slice preparations, a series of 20 doughnuts shifted orthogonally to the in-plane XY diagonal was designed to match the soma dimensions of the target cell, with each doughnut scanned for 70 µs, yielding a total single-cell excitation duration of 1.4 ms **(Figure 7C, left)**. Representative photocurrents recorded in whole-cell voltage clamp from a single MLI under concurrent two-photon photostimulation at increasing laser powers are shown in **Figure 7C**. Notably, photocurrents were initiated before the full sequence of 20 doughnuts had been delivered, confirming the efficiency of this scanned doughnut strategy for sub-millisecond photoexcitation.

#### Photocurrent characterization and spatial resolution

The input–output relationship between photocurrent amplitude and laser power followed a saturation curve with a half-saturation power √P_0_^2^ of 8 mW **(Figure 7D)**, indicating that substantial photocurrents can be evoked at low applied power. The spatial specificity of two-photon doughnut photostimulation was characterized by measuring photocurrent amplitude as a function of lateral and axial displacement of the ULoVE focus relative to the MLI soma **(Figure 7E)**. The lateral resolution constant corresponded to 0.72 cell diameters, and the axial resolution was 8.3 µm — both below the typical MLI soma diameter of 9–10 µm — confirming that individual somata can be selectively photostimulated without co-activating neighboring cells. **Sequential multi-cell photostimulation and functional connectivity mapping**. The sub-millisecond single-cell excitation duration and cellular-scale spatial resolution together enable sequential photostimulation of multiple individual MLIs within a single revisit cycle. To demonstrate this capability and exploit the cell-type specificity of the Gateway-cloned expression constructs, nine MLIs in the vicinity of a patch-clamped Purkinje cell were sequentially photostimulated at a revisit cycle frequency of 710 Hz, with each MLI receiving a 1.4 ms multi-doughnut excitation per cycle **(Figure 7F)**. Inhibitory postsynaptic currents (IPSCs) were recorded at the Purkinje cell soma to identify which of the nine photostimulated MLIs were presynaptically connected. Of the nine cells probed, only MLI #8 generated a reliable postsynaptic response, producing an average IPSC of 46 pA across 10 consecutive trials with one failure **(Figure 7G)**. The remaining eight photostimulated MLIs produced no detectable postsynaptic current, confirming the specificity of the photostimulation and, interestingly, showing a sparse nature of MLI-to-Purkinje cell connectivity at the single-synapse level.

Taken together, these results demonstrate that the combination of Gateway-cloned cell-type-specific viral vectors and ULoVE doughnut photostimulation enables functional optogenetic interrogation of inhibitory microcircuits with single-cell resolution and sub-millisecond temporal precision. The ability to probe synaptic connectivity across multiple candidate presynaptic cells within a single recording — made possible here by the 710 Hz revisit rate — illustrates how the Gateway pipeline’s capacity to generate precisely targeted, density-controlled expression constructs directly enables advanced optical interrogation strategies that would otherwise be technically inaccessible.

## Discussion

We have established a modular Gateway-based cloning pipeline for the rapid construction of rAAV expression vectors tailored to optical neuroscience applications. Gateway cloning has previously been applied to rAAV vector construction in neuroscience, including a molecular toolbox for up- and down-regulation of neuronal gene expression (White and Nolan, 2011) and for targeted viral vector transduction using novel cell-type-specific promoters (Wykes et al., 2019). Our approach builds on this foundation by explicitly exploiting the combinatorial modularity of the system as a core design principle: by maintaining organized libraries of promoter and transgene entry clones that can be recombined in any combination through a single overnight LR reaction, the pipeline substantially reduces the time and effort required to generate, optimize, and diversify viral expression constructs at scale. The results presented here demonstrate that this pipeline is not merely a technical convenience but an enabling platform: the cell-type-specific, density-tunable, and multi-indicator expression strategies it supports were essential preconditions for the ULoVE-based optical recordings, optogenetic and histology-tracing experiments described above, which would have been prohibitively time-consuming with conventional cloning approaches. Equally, the rAAV production protocol described here — based on simple polyethylenimine (PEI)-mediated (Longo et al., 2013) triple transfection of HEK293T cells followed by iodixanol gradient purification — is accessible to any laboratory with standard cell culture infrastructure and an ultracentrifuge, and routinely yields titers of 10^13^–10^14^ GC/ml, which are sufficient for in vivo use. Combining a streamlined cloning pipeline with in-house production capability removes two principal bottlenecks in optical neuroscience tool development, enabling generation of injection-ready viral vectors from a new transgene sequence in as little as two to three weeks.

When considering the design of optical tools within this framework, several points warrant attention. Packaging capacity management is an important constraint: the AAV packaging limit of approximately 4.7 kb, including ITRs, presents challenges for neuroscience vectors, which often contain relatively large coding sequences (GECI variants ∼1.3 kb, ChR2-EYFP ∼1.8 kb, GEVI plus targeting motifs ∼1.5 kb) combined with regulatory and promoter elements. Gateway cloning facilitates rapid assessment of different configurations to maximize functional cargo while respecting packaging constraints (Wu et al., 2010). Fluorescent protein fusion optimization is another consideration: many optogenetic actuators and indicators incorporate fluorescent protein domains for visualization, either fused to the gene-of-interest or co-expressed via P2A ribosomal-skipping sequences (Lo et al., 2015; Liu et al., 2017; Rao et al., 2025). Gateway-based modular design enables systematic testing of different linker sequences and fluorescent protein variants as they become available and characterized (Shaner et al., 2005; Lambert et al., 2019). Subcellular targeting is also modularly addressable: membrane-targeting signals, nuclear import sequences for recombinases, and other subcellular trafficking motifs such as Golgi transit and ER export signals (Gradinaru et al., 2010) can be incorporated through Gibson assembly (see Methods section) and implemented into rAAVs by Gateway cloning.

Regarding neuronal GEVI recordings specifically, we and others have found that incorporating the soma localization motif from Kv2.1 (Lim et al., 2000; Jensen et al., 2017; Lim and Liu, 2018) at the C-terminus of the GEVI is of particular utility for reducing neuropil background signal and facilitating high signal-to-noise recordings (Villette et al., 2019; Lee et al., 2022). On the other hand, we have found that GEVI expression in non-neuronal cells such as astrocytes is impaired when the Kv2.1 soma targeting motif is present (unpublished), and that despite using this motif for GEVI expression in cerebellar Purkinje cells, we observe extensive signal in the dendritic tree (**see Figure 5**), further illustrating the value of the flexibility afforded by the Gateway pipeline for evaluating expression constructs across cell types. A further consideration inherent to Gateway-based cloning is that the recombination reaction leaves behind short (21–25 bp) “attB scar” sequences flanking each assembled element in the final pEXPR construct. We and others have observed no evidence that these residual sequences impair transgene expression, transcript stability, or AAV packaging in any of the constructs described here, consistent with reports that attB sites do not detectably alter transcription kinetics, expression pattern, or mRNA stability in other Gateway-based expression systems (Perehinec et al., 2007; Giuraniuc et al., 2013), though Perehinec et al. (2007) noted a modest reduction in absolute expression level when att sites were positioned immediately adjacent to a promoter or ribosome-binding site.

MultiSite LR Clonase reactions require some additional care relative to standard two-fragment reactions. The simultaneous assembly of three components demands precise stoichiometric balancing of entry clones — typically at equimolar ratios, with empirical adjustment where one component is rate-limiting — and high-quality supercoiled plasmid preparations, as nicked or degraded DNA significantly reduces efficiency when three recombination events must occur concurrently. Extended incubation times (24–48 hours, with optional addition of fresh LR Clonase Plus enzyme at the midpoint) and the use of LR Clonase Plus enzyme formulated specifically for MultiSite reactions both improve yields. Because MultiSite reactions can generate more background colonies from incomplete assemblies than standard two-fragment reactions, more rigorous colony screening is warranted: junction PCR spanning each fragment boundary, diagnostic restriction digestion confirming all elements are present, and full sequencing of the assembled expression cassette are recommended before proceeding to AAV production. Complete details of stoichiometric ratios, reaction volumes, and screening protocols for specific construct types in our library are provided in the Materials and Methods.

The destination vector framework also accommodates self-complementary AAV (scAAV) configurations, generated by deletion of the terminal resolution site (trs) from one ITR, which accelerate the onset of transgene expression by bypassing the requirement for second-strand synthesis in transduced cells (McCarty et al., 1994, 2003). Although not employed in the current study, this option can be engineered into the existing pDEST plasmids and may be useful for applications requiring rapid transgene onset, with the caveat that packaging capacity is halved and ddPCR quantification requires adjusted protocols to account for the unusual secondary structures of the trs-deleted ITR (Wilmott et al., 2019).

The Purkinje cell recordings presented in this study illustrate these principles concretely. The ability to rapidly achieve matched, cell-type-specific expression across three indicator classes --- two calcium sensors (GCaMP6f and jRGECO1a) and a soma-targeted voltage indicator (JEDI2P-Kv) --- within a single experimental series, using a common Gateway-constructed L7::Cre driver virus to conditionally express each indicator and to tune expression density over two orders of magnitude, was essential for establishing the ULoVE recording conditions. Resolving dendritic spikelets with a genetically encoded voltage indicator at ∼5 kHz in awake animals, previously reported using intracellular electrophysiology (Maruta et al., 2007; Kitamura and Häusser, 2011) and the synthetic voltage dye ANNINE-6+ (Roome and Kuhn, 2018), was achieved here by combining the sparse, cell-type-specific expression enabled by Gateway-built driver and indicator vectors with the high photon-flux and motion-resilient excitation provided by ULoVE. Moreover, the comparative kinetic analysis across indicator classes — revealing that climbing fiber-evoked calcium transients in Purkinje dendrites rise substantially faster than predicted from cortical benchmarks, yet still 6.7-fold slower than the underlying voltage depolarization — was facilitated by the Gateway pipeline, which made it practical to generate, validate, and deploy matched driver viruses and voltage-indicator vectors in parallel. This type of systematic multi-indicator comparison, which would be prohibitively time-consuming with conventional cloning, exemplifies the experimental leverage that modular vector construction provides.

## Conclusions

Gateway cloning provides a powerful solution to a recurring challenge in optical neuroscience: the need to rapidly generate, compare, and optimize rAAV expression constructs across multiple promoters, transgene sequences, and conditional expression configurations. By maintaining modular libraries of sequence-verified entry clones that can be assembled into functional expression plasmids in a single overnight reaction, the Gateway pipeline substantially compresses the design-build-test cycle that underpins iterative tool development. The 105 viral productions described here, spanning 9 promoter elements and more than 20 transgene configurations, illustrate the practical scale at which this combinatorial leverage operates.

The integration of this pipeline with advanced optical acquisition methods underscores a broader principle: the value of a molecular tool development platform is determined not only by what it can build in isolation, but also by what it makes possible experimentally. As optical tools continue to proliferate — with new calcium indicators, voltage sensors, and optogenetic actuators emerging at an accelerating pace — the ability to rapidly incorporate and systematically compare new transgene entries within an existing library framework ensures that adoption of these advances remains efficient and reproducible across experimental contexts.

Looking forward, the modular architecture of the Gateway pipeline positions it well for emerging applications in all-optical interrogation and closed-loop circuit control, where simultaneous expression of multiple actuators and reporters in defined cell populations will be increasingly required. The ability to construct such multi-component vectors rapidly and in parallel — rather than through sequential bespoke cloning — will be an important practical advantage as experimental designs grow in complexity.

The success of Gateway cloning for neuroscience AAV vectors ultimately reflects a broader principle: tool development in biological research benefits enormously from standardized, modular approaches that minimize technical overhead and maximize experimental flexibility. As neuroscience increasingly relies on sophisticated genetic tools for circuit interrogation, Gateway cloning stands as an essential enabling technology bridging molecular biology and systems neuroscience.

The Purkinje cell dendritic recordings and MLI optogenetic experiments presented here serve as proof of concept for this integrated approach. The Gateway cloning pipeline provided the expression toolkit; ULoVE provided the optical resolution. Neither alone would have been sufficient — sparse, cell-type-specific expression enabled by Gateway was a prerequisite for the high signal-to-noise ULoVE recordings, while ULoVE’s temporal resolution revealed biology that slower acquisition methods would have missed entirely. Together, they represent a blueprint for combining modular molecular tool development with advanced optical acquisition — an approach that generalizes to any experimental system where cell-type specificity, expression density control, and high-speed optical recording must be optimized simultaneously. As the optical toolbox continues to expand, the Gateway pipeline ensures that adoption of these advances remains rapid, systematic, and reproducible.

## Material and Methods

### Gateway cloning

#### pENTR plasmid construction

The initial phase involves the generation of pENTR plasmids containing optical tool coding sequences or promoters for general neuronal or cell-type-specific expression. Optical tool coding sequences (e.g., Chronos, JEDI2P) are amplified using a high-fidelity polymerase with primers (specific primer sequences can be made available on request) that flank the GoI with attB1 and attB2 BP-recombinase sequences (Hartley et al., 2000). Primer design generally accounts for a Kozak sequence to ensure optimal translation initiation (Kozak, 1999). The attB-flanked PCR products undergo BP recombination with pDONR vectors, typically achieving 85– 95% recombination efficiency under optimized conditions, producing the GoI flanked by attL recombination sites. For conditional GoI expression (CRE or FLPo-dependent), transgenes were instead inserted by Gibson assembly into a pre-configured pENTR plasmid in an inverted conditional configuration, as described below (see Transgene entry clone library; **Supplemental Figure 2**). Following BP recombination or Gibson assembly, pENTR clones underwent comprehensive Sanger sequencing to confirm the absence of polymerase-introduced mutations. Sequence-verified pENTR clones serve as permanent repositories, eliminating the need for repeated amplification and reducing mutation accumulation. Primer design and *in silico* reactions were performed using SnapGene software (www.snapgene.com)

#### LR recombination and vector assembly

LR reactions transfer promoter sequences and/or coding sequences from pENTR plasmids into AAV pDEST plasmids. In our hands, LR reactions for AAV vectors are routinely performed for 1.5 hours @25°C. Following transformation and after both positive (antibiotic) and negative genetic selection (ccdB), colonies are screened by diagnostic restriction digestion with SmaI to confirm ITR sequence integrity, and internal restriction digestion of coding sequences before sequencing. For neuroscience applications where vector functionality is critical, selected clones should undergo complete sequencing of the expression cassette, including promoter, coding sequence, and polyadenylation signal. Transfection of HEK293, CHO, or neuronal cell lines provides in vitro assessment of expression levels and protein localization before rAAV production. For optogenetic vectors, patch-clamp electrophysiology confirms photocurrent generation; for calcium and voltage indicators, stimulus-evoked fluorescence responses validate sensor functionality (Tian et al., 2009; Villette et al., 2019). AAV ITRs are susceptible to recombination and deletion during bacterial propagation; we utilize a low-copy-number origin of replication on all ITR-containing plasmids, which in our hands has reduced the occurrence of ITR mutations.

#### Construction of pENTR and pEXPR plasmids — worked example

pAAV(2655).TRE.fDiO.JEDI-2P-Kv (pEXPR A-193). The construction of pEXPR A-193 illustrates the three-stage Gateway cloning workflow used to generate conditional rAAV expression constructs in this study. All entry clones generated were sequence-verified by Sanger sequencing across the full expression cassette prior to use and stored as bacterial glycerol stocks at −80°C.

##### Stage 1 — BP recombination

*construction of pENTR-L1-TRE-R5 (A-105)*. The TRE promoter sequence was amplified by high-fidelity PCR from pAAV.TRE.EYFP.WPRE (Addgene #104111, A-99) using Phusion HF polymerase (Thermo Fisher) with primers appending attB1 and attB5r recombination sequences at the 5′ and 3′ ends, respectively (forward primer AA137, reverse primer AA138). Each 25 µl reaction contained 1× Phusion Green HF Buffer, 0.2 mM dNTPs, 0.5 µM each primer, 2 ng template DNA, and 0.02 U/µl Phusion polymerase. Thermal cycling: 98°C 3 min; 30 cycles of 98°C 30 sec, gradient annealing 50–70°C 30 sec, 72°C 25 sec; final extension 72°C 10 min. The amplified TRE fragment (387 bp) was purified by agarose gel extraction and NucleoSpin Gel/PCR Clean-Up, eluted in 30 µl NE buffer at 70°C, and quantified by NanoDrop. BP recombination was performed by combining 0.05 pmol attB-flanked PCR product (12.6 ng) with 0.05 pmol pDONR221 P1-P5r (A-2; 154.9 ng) in TE buffer pH 8.0 to a total volume of 8 µl, adding 2 µl BP Clonase II (Thermo Fisher, stored at −80°C), and incubating at 25°C for 1 hour followed by 1.5 hours at 25°C. Proteinase K solution (1 µl) was added, and the reaction was incubated at 37°C for 10 minutes to terminate. 2.5 µl of the BP reaction was transformed into 25 µl DH5α competent cells (NEB C2987H) by heat shock at 42°C for 30 seconds, 5 min on ice, followed by addition of 475 µl SOC, followed by addition of 475 µl SOC at room temperature, and incubated at 37°C for 1 hour at 220 rpm before plating on LB-kanamycin agar. Minipreps were screened by diagnostic restriction digestion (BspHI for vector linearization; BstYI for internal coding verification). Clone #BP TRE3 (pENTR-L1-TRE-R5, A-105; 87.5 ng/µl, A260/A280 1.9) was selected for use in subsequent LR reactions.

##### Stage 2 — Gibson assembly

###### construction of pENTR-fDiO-JEDI-2P-Kv (A-148)

The JEDI-2P-Kv coding sequence (1555 bp, including the Kv2.1 soma-targeting motif) was amplified from pAAV.eF1α.DiO.JEDI-2P-Kv (A-143, gift from Francois St-Pierre, Baylor College of Medicine) using Phusion HF polymerase with primers incorporating overlapping sequences matching the fDiO insertion site of the linearized recipient vector (forward primer AA59/AA23, reverse primer Fragment.REV). Thermal cycling: 98°C 3 min; 34 cycles of 98°C 10 s, 50–65°C 30 s, 72°C 47 s; final extension 72°C 10 min. The 1555 bp fragment was purified by agarose gel extraction (eluted in 30 µl NE buffer at 70°C; 157.95 ng/µl). Gibson assembly was performed using <u>New England Biolabs (NEB) Gibson Assembly Cloning kit</u> (M5510A), combining 0.04 pmol JEDI-2P-Kv PCR insert (40.4 ng) with 0.02 pmol linearized pUC57 two-fragment Gateway fDiO vector (39.7 ng) in a total volume of 20 µl in 1× Gibson Assembly Mix, and incubating at 50°C for 20 minutes. We prepared the linearized vector as a batch stock by inverse PCR of the pUC57 fDiO plasmid using primers Gibson pUC57 Flex2-F (5’-CATGGTGGCGCTAGGGGCCCGCGGTACCG-3’) and Gibson pUC57 Flex2-R (5’-TAAGGGGATCCTAGTGGCGCGCC-3’), gel-purified it, and stored it at −20°C for repeated use across multiple Gibson assembly reactions. Both the frt/fDiO and lox/Flex pUC57 backbone plasmids were designed to be linearizable with the same primer pair, enabling a single-batch preparation protocol to support all conditional transgene-entry clone constructions regardless of the recombinase system. Both plasmids were custom-synthesized by GeneCust, Luxembourg (pUC57 L5 Atasoy Flex switch L2 and pUC57 L5 Fenno fDiO L2, respectively). 1.5 µl (6 ng) was transformed into 25 µl DH5α competent cells, recovered in 475 µl SOC, and plated on LB-kanamycin agar. Minipreps were screened by Sanger sequencing using primers KP GFP1, AA119, and AA121. Clone #G12 (pENTR-fDiO-JEDI-2P-Kv, A-148) was confirmed by complete sequence verification and stored as a bacterial glycerol stock.

##### Stage 3 — MultiSite LR recombination

###### construction of pAAV(2655).TRE.fDiO.JEDI-2P-Kv (A-193)

The final pEXPR plasmid was assembled by MultiSite LR recombination using Gateway LR Clonase II Enzyme Mix (Thermo Fisher, ref. 11791020). Each reaction combined three components at equimolar amounts (0.04 pmol each): pENTR-L1-TRE-R5 (A-105; 0.9 µl, 76.4 ng), pENTR-fDiO-JEDI-2P-Kv (A-148 #G12; 2.5 µl, 118.9 ng), and pDEST-SMD2-Rfa-WPRE (A-63 clone 3, freshly prepared from glycerol stock; 0.4 µl, 174.7 ng), made up to 8 µl with TE buffer pH 8.0 and supplemented with 2 µl LR Clonase II, for a total reaction volume of 10 µl. The reaction was incubated at 25°C for 1.5 hours, terminated with 1 µl Proteinase K solution at 37°C for 10 minutes, and 0.3 µl (13.5 ng) was transformed into 25 µl DH5α competent cells (NEB C2987H) by heat shock, recovered in SOC, and plated on LB-ampicillin agar. Minipreps from 14 colonies were screened by diagnostic restriction digestion with MscI and SmaI. Clones #5, #6, #10, and #14 passed both digestion screens; clone #14 was selected for complete Sanger sequencing of the expression cassette, which confirmed the correct assembly. pAAV(2655).TRE.fDiO.JEDI-2P-Kv (pEXPR A-193, clone #14) was stored as a bacterial glycerol stock at −80°C and used directly for rAAV production.

### Design and library construction

#### Promoter entry clone library

Promoter elements were cloned into pENTR plasmids flanked by attL1 and attR5 recombination sites, suitable for use as the promoter component in MultiSite LR reactions. The library currently encompasses promoters for ubiquitous AAV expression (CAG, CMV, eF1α, PGK), pan-neuronal expression (hSyn, CaMKIIα), astrocyte-specific expression (GfaABC1D GfaABC1D-βG), and intersectional strategies using a tetracycline-responsive element (TRE) in combination with a conditionally expressed tetracycline transactivator (tTA). Three fragments of the L7/pcp2 promoter (0.6kb (L7_0·6_), 1kb (L71), and 3kb (L7_3_)) (Nitta et al., 2017) and a 2.2 kb fragment of the c-kit promoter (Villette et al., 2019) were also cloned to enable Purkinje cell-specific and molecular layer interneuron-specific expression, respectively.

#### Transgene entry clone library

Transgene sequences were cloned into pENTR plasmids flanked by attL5 and attL2 recombination sites. For unconditional expression, sequences were cloned directly following a Kozak consensus sequence. For conditional (CRE- or FLPo-dependent) expression, transgenes were inserted by Gibson assembly into a pre-configured pENTR plasmid carrying heterotypic pairs of lox or frt recombination sites in an inverted configuration, such that recombinase activity catalyzes permanent reversion of the transgene to its functional orientation (Atasoy et al., 2008; Fenno et al., 2014); **Figure 3, Supplemental Figure 2**. The transgene library currently includes, but is expanding rapidly, (see Supplemental **Table 1**) optogenetic actuators (Chronos, ChrimsonR, iC++, PsChR2), voltage indicators (ASAP3b-Kv, JEDI-2P-Kv, JEDI-2P-TS-ER-Kv2.1PRC, VADER1, FORCE1f-Kv), fluorescent reporters (tdTomato, mRuby2, eGFP, eGFP-CAAX, iRFP670, Crimson-CAAX), recombinases (CRE, nls-FLPo), transcriptional activators (tTA), and other GoIs such as the biotin ligase BirA fused to tau for proximal biotinylating (tau-BirA) or the mouse canine adenovirus (CAV) receptor (mCAR), whose expression allows retrograde CAV transduction.

#### Destination vectors

All rAAV expression constructs were generated using destination vectors (pDEST) containing AAV2 inverted terminal repeats (ITRs) flanking attR1 and attR2 recombination sites, along with a ccdB-CmR negative selection cassette. The ITRs within destination vectors must maintain their palindromic structure and specific spacing to ensure efficient replication and packaging of the resultant rAAV (Grimm et al., 1998). When the plasmid origin of replication is high-copy (i.e., pUC-based), the ITR adjacent to the origin is frequently found to carry an 11 bp deletion in the B or C domain, thought to be repaired during the in vitro replication and second-strand synthesis phase of rAAV production (Samulski et al., 1983; Feiner et al., 2019; Shitik et al., 2023; Chen et al., 2024). Although whether this deletion ultimately affects viral vector production remains unclear, we prefer to use destination vectors with a low-copy-number origin (i.e., pBR-based) that maintain fully intact ITRs, and to propagate all ITR-containing plasmids in DH5α bacteria to minimize ITR instability. In addition to the ITR-flanked expression cassette, pDEST vectors incorporate post-transcriptional control elements to enhance transgene expression: a Woodchuck Hepatitis Virus post-transcriptional regulatory element (WPRE) to enhance mRNA stability and transgene expression levels (Loeb et al., 1999), and a bovine growth hormone polyadenylation signal (bGH-pA) for efficient mRNA maturation. Together, these elements total 862 bp in the standard destination vector backbone (A-63, pDEST.pAAV(2566)SMD2.Rfa.WPRE.BGHpA), which was used for the majority of constructs reported in this study. For constructs approaching the AAV packaging limit of approximately 4.7 kb, a second destination vector backbone was used (A-186, pDEST.pAAV(2566)SMD2.Rfa.W3SL.SV40pA), in which minimized versions of these post-transcriptional elements — WPRE3 in place of WPRE, and SV40 late-pA in place of bGH-pA (Choi et al., 2014; Li et al., 2018) — reduce the combined regulatory element footprint to 356 bp, freeing approximately 500 bp of additional packaging space for larger transgene or promoter sequences. For a pDEST vector to have maximal versatility, it should not contain residual control regions that could influence the promoter used to drive the GoI or the function of the GoI itself. White and Nolan (2011) introduced a pDEST plasmid (pAAV-Gateway, Addgene #32671) as part of a molecular toolbox for rapid viral vector generation, but subsequent characterization indicated it harbors remnants of the CAG promoter from its parent plasmid pCAGW-ChR2-Venus (Petreanu et al., 2009); Matthew Nolan, personal communication), which could confound experiments using cell-type-specific or weaker promoters. Our destination vectors were designed and verified to be free of such residual regulatory elements. Upon MultiSite LR recombination with the appropriate promoter and transgene entry clones, the ccdB-CmR cassette is replaced by the assembled expression cassette, yielding a functional pEXPR plasmid ready for rAAV production (**Figure 3**).

### rAAV production and purification

#### Cell culture and expansion

rAAV vectors were produced by triple transfection of HEK293T cells followed by iodixanol gradient purification, using a protocol adapted from (Challis et al., 2019). HEK293T cells (ATCC®, CRL-3216™) were maintained in Dulbecco’s Modified Eagle Medium (DMEM) supplemented with 10% fetal bovine serum (FBS) and expanded under standard conditions (37°C, 5% CO_2_). For each production, cells were expanded progressively from a single cryopreserved vial (4.2×10^6^ cells) through a series of passages: initial recovery in a 10 cm dish (48hrs), followed by sequential expansion into four 10 cm dishes (1:5 for 72hrs), then into four T150 flasks (48hrs), and finally into twenty T150 flasks at a 1:4 split ratio. On the day of transfection (72 hrs after plating), medium was replaced with 15 ml fresh DMEM per T150 flask 4–5 hours before the transfection mixture was applied. Cells were required to be 80– 90% confluent at the time of transfection.

#### Triple transfection

Five T150 flasks (150 cm^2^ growth area each) were used for triple transfection with PEI (Polysciences, Inc., Eppelheim, Germany; 1 mg/ml in water) as the transfection reagent. Each flask received a total of 28 µg of plasmid DNA distributed across three plasmids: 12 µg helper plasmid (pAdDeltaF6, Addgene #112867), 10 µg AAV capsid plasmid, and 6 µg of the Gateway-generated pEXPR vector containing the transgene of interest flanked by AAV2 ITRs. As the ITR type was constant across all constructs, rAAV vectors are identified throughout by capsid type alone (e.g., AAV1, AAV5), rather than the conventional AAV2/x notation. DNA concentrations were adjusted to a minimum of 1.0 µg/µl before use. For each production, all three plasmid solutions were combined in 2.5 mL DMEM (without serum) in a 15 mL conical tube and mixed thoroughly. PEI was then added at a PEI:DNA mass ratio of 4.2:1. The mixture was vortexed for 10 seconds and incubated at room temperature for 5–10 minutes before the calculated volume was added to each flask and distributed by gentle rocking.

Approximately 12–16 hours post-transfection, 5 mL of DMEM supplemented with 10% FBS was added to each flask without replacing the existing medium. The volume of transfection mixture added per flask was calculated from the individual plasmid concentrations using a standardized transfection spreadsheet (Supplemental AAV production worksheet), targeting a total DNA mass of 28 µg per flask at a DNA mass ratio of 2:1.67:1 (helper:capsid:transgene), or 2:4:1 when using any non-natural/engineered capsid type.

#### Harvest

AAV particles were harvested 120 hours post-transfection via two parallel streams — conditioned medium and cell lysate — which were processed separately before being combined. Earlier harvests at 72 hours were found to yield lower titers and were therefore discontinued in favor of the single 120-hour harvest.

##### Cell lysate stream

Cells were scraped from the flask surface and collected together with the conditioned medium by centrifugation at 2,000 × g for 10 minutes at 4°C. The cell pellet was resuspended in 10 ml lysis buffer (50 mM Tris-Cl pH 8.5, 150 mM NaCl) and subjected to three freeze-thaw cycles using a dry ice/ethanol bath and a 37°C water bath, with vortexing between each cycle (minimum 15 minutes per freeze). The lysate was supplemented with MgCl_2_ to a final concentration of 10 mM, and DNase I was added to 0.1 mg/ml final concentration (110 µl of a 10 mg/ml stock; Roche 11284932001). After 30 minutes of incubation at 37°C, the lysate was clarified by centrifugation at 3,700 × g for 20 minutes at 4°C, and the supernatant was retained.

##### Conditioned medium stream

The supernatant from the initial 2,000 × g centrifugation was supplemented with 5× polyethylene glycol (PEG) solution (40% PEG w/v, 2.5 M NaCl; Sigma 89510) to a 1× final concentration (26.5 mL per 105 mL medium) and incubated at 4°C for a minimum of 2 hours and up to 3 days. The resulting precipitate was collected by centrifugation at 3,700 × g for 20 minutes at 4°C. The pellet was resuspended in the clarified cell lysate, and the combined suspension was kept overnight at 4°C (or at 37°C for 1 hour) to facilitate complete resuspension, which can be maintained for up to 7 days at 4°C prior to gradient purification.

### Iodixanol gradient ultracentrifugation

The combined lysate/PEG resuspension was loaded into a Beckman QuickSeal ultracentrifuge tube (344623) and underlaid with sequential iodixanol density gradient layers (OptiPrep; Sigma D1556) in the following order from top to bottom: 11 ml lysate, 9 ml 15% iodixanol, 5 ml 25% iodixanol (supplemented with 100 µl 0.5% phenol red for interface visualization; Sigma P-0290), 5 ml 40% iodixanol, and 3 ml 60% iodixanol (supplemented with 100 µl phenol red). The tube was filled to the base of the neck with lysis buffer if necessary. Iodixanol solutions were prepared in 5× PBS-MK (250 ml PBS 10×, 2.5 ml 1 M MgCl_2_, 6.25 ml 1 M KCl, water to 500 ml) with 5 M NaCl as indicated. Gradients were centrifuged in a Beckman 70Ti rotor at 350,000 × g (59,000 rpm) for 120 minutes at 20°C.

Following centrifugation, an 18G needle was inserted at the top of the gradient tube to allow air entry, and the 40% iodixanol fraction (containing rAAV particles) was withdrawn from the side of the tube using a 10 mL syringe pre-loaded with 4 mL of PBS/0.001% Pluronic F-68. The fraction was withdrawn slowly and without interruption to avoid disturbing the 25%/40% interface. If interface contamination was suspected, the fraction was pre-filtered through a 0.45 µm syringe filter before the subsequent 0.22 µm filtration step.

### Concentration and buffer exchange

The recovered 40% fraction and pre-loaded buffer were passed through a pre-washed 0.22 µm syringe filter directly into an Amicon Ultra-15 centrifugal filter unit (Ultracel-100 membrane, Millipore PL100) that had been pre-conditioned by sequential washes with PBS/0.1%, 0.01%, and 0.001% Pluronic F-68, with centrifugation at 4,000 × g for 5 minutes between each wash. The loaded Amicon device was centrifuged at 4,121 × g for 20 minutes until the volume was reduced to approximately 500 µl. The retentate was then washed three additional times with 5–7 mL PBS/0.001% Pluronic F-68, with centrifugation between each wash, and the final volume was reduced to 100–250 µl during the last spin. The purified rAAV was recovered from the filter membrane and stored at 4°C for up to one month, or aliquoted on dry ice and stored at –80°C for long-term use.

### Reagents and solutions

Lysis buffer (500 mL): 4.38 g NaCl (150 mM), 25 mL 1 M Tris-Cl pH 8.5 (50 mM final), autoclaved. 5× PEG solution: 400 g PEG (Sigma 89510), 146.25 g NaCl, water to 1 L, sterile-filtered through a 0.22 µm Steritop filter. Iodixanol fractions prepared as follows (per 200 ml): 15% — 50 ml iodixanol, 40 ml 5 M NaCl, 40 ml 5× PBS-MK, 70 ml water; 25% — 83.2 ml iodixanol, 40 ml 5× PBS-MK, 76.8 ml water, 100 µl phenol red; 40% — 133.2 ml iodixanol, 40 ml 5× PBS-MK, 26.8 ml water; 60% — 200 ml iodixanol, 100 µl phenol red.

### Titer determination

Genome copy titers of purified rAAV preparations were determined by droplet digital PCR (ddPCR). Results are expressed as genome copies per milliliter (GC/ml). Because physical titer does not always correlate with functional transduction efficiency, and because, in our hands, ddPCR measurements of the same rAAV preparation performed in different laboratories can differ by up to fourfold, initial pilot injections were performed across a range of titers to establish optimal experimental conditions before large-scale use.

### Genome copy titer determination by droplet digital PCR

Primers targeted the AAV2 inverted terminal repeat (ITR) sequence, and ddPCR followed the protocols described in (Aurnhammer et al., 2012; Lock et al., 2014).

#### Sample preparation

Prior to ddPCR, rAAV samples were first treated to degrade unencapsidated DNA and render the capsid accessible for protease digestion. One microliter of purified virus was diluted in DNase I reaction buffer (1x: 20 mM Tris-Cl pH 8.3, 2 mM MgCl_2_; Sigma AMP-D1) to a total volume of 50 µl containing 1 unit DNase I, and incubated for 30 minutes at 37°C. From 2021 onwards, to minimize non-specific adsorption during serial dilution, samples were supplemented with 0.05% Pluronic F-68 (Gibco, 24040-032). DNase I was inactivated by heating to 65°C for 10 minutes. Proteinase K (10 µg; Sigma P2308, 20 mg/ml stock in PCR-grade water) was then added to the same 50 µl volume and incubated for 60 minutes at 50°C to digest capsid proteins and release encapsidated viral genomes. Proteinase K was inactivated by heating to 95°C for 20 minutes. Serial dilutions of each sample were prepared in 1× dilution buffer (1× GeneAmp PCR Buffer I, Applied Biosystems; 0.05% Pluronic F-68; PCR-grade water) across dilutions of 20,000× and 180,000×. Dilutions spanning the linear quantification range of the assay were selected for each sample based on preliminary assessment, with the rAAV8 reference standard (ATCC®, VR-1816) included in each run for both inter-run normalization and external normalization according to the expected value for qPCR for rAAV8 from the consortium AAV Reference Standard Working

#### ddPCR reaction

Each 25 µl ddPCR reaction contained: 5 µl Perfecta Multiplex qScript ToughMix (5×, used at 1× final concentration), 2.5 µl fluorescein (1 µM stock, 100 nM final), 4.1 µl primer/probe mix (25×, used at 1× final), 5 µl diluted DNA template, and 8.4 µl PCR-grade water. The ITR-targeting primer and probe sequences were: forward primer 5’-GGAACCCCTAGTGATGGAGTT-3’ (0.72 µM final); reverse primer 5’-CGGCCTCAGTGAGCGA-3’ (0.72 µM final); FAM-labelled probe 5’-CACTCCCTCTCTGCGCGCTCG-3’ (0.2 µM final; dual-labelled with FAM at 5’ and BBQ650 quencher at 3’; Eurofins, HPLC-purified). Reactions were partitioned into 30,000 droplets using Sapphire chips (4 samples per chip, 3 chips per run) on the Naica™ Crystal droplet digital PCR system (Stilla Technologies). Thermal cycling conditions were: 95°C for 10 minutes (initial denaturation); 55 cycles of 95°C for 30 seconds (denaturation) and 60°C for 1 minute (annealing/extension). Results are expressed as genome copies per milliliter (GC/ml), calculated from the cdPCR absolute quantification output corrected for the dilution factor applied and the expected VR-1816 control value.

### Stereotaxic viral injections for anterograde tracing of DCN projections (Figure 4) and validation of cell-type-specific rAAV drivers (Figure 5)

All protocols adhered to the French National Ethics Committee for Sciences and Health report on Ethical Principles for Animal Experimentation and the European Community Directive 86/609/EEC, under agreements #29791 and #57060.Group (AAVRSWG).

Stereotaxic injections were performed on adult vGlut2::CRE (Borgius et al., 2010); GlyT2::eGFP (Zeilhofer et al., 2005) double-transgenic mice (males, injected at approximately 4 months of age). Anesthesia was induced by intraperitoneal injection of a ketamine/xylazine mixture. A working solution was prepared from stock ketamine (Ketamine 1000, 10 mg/ml) and xylazine (2% Rompun, 20 mg/ml) by diluting 0.37 ml ketamine and 0.25 ml xylazine into 10 ml saline; mice received 37 µl of diluted stock per gram body weight, corresponding to 14.8 µg ketamine/g and 20 µg xylazine/g. Two-thirds of the anesthetic dose was administered first, and depth of anesthesia was confirmed by loss of the foot-retraction reflex before the remaining third was administered. The animal was mounted in a stereotaxic frame with the tongue clear of the airway, ophthalmic balm was applied to protect the eyes, and the eyes were shaded from the surgical lamp.

A longitudinal midline incision was made from behind the eyes to the caudal extent of the skull to expose the parietal and interparietal bones, and the skull was cleared of overlying tissue. The skull was levelled by matching dorsoventral (z) measurements at Bregma and lambda. Injection coordinates were defined relative to Bregma: x = 0.6 mm (mediolateral) and y = −6.5 mm (anteroposterior), targeting the medial DCN. A 0.5 mm burr hole was drilled at the injection site, stopping just before full penetration, and the remaining thin bone was carefully removed with a 27G needle. The dorsoventral coordinate (z) was measured from the surface of the exposed brain at the injection site.

A quartz glass capillary pipette, back-filled with mineral oil and mounted on a hydraulic injection system, was used for all injections. The pipette was loaded with the viral mixture, positive pressure was applied to confirm flow at the tip, and the pipette was lowered to a depth of 2.45 mm below the brain surface and, after a 2-minute wait, retracted by 50 µm to 2.4 mm to create a small reservoir at the target. A total volume of 150 nl was injected at a rate of 100 nl/min. The pipette was left in place for 10 minutes after injection before being slowly retracted. The incision was closed with a running suture, the animal received 0.4 ml saline subcutaneously, and was allowed to recover under a heat source (<33°C) for approximately 20 minutes.

For sparse anterograde labelling of vGlut2+ DCN projection neurons **(Figure 4)**, three Gateway-cloned or commercially sourced AAVs were co-injected: AAV1.eF1α.DiO.nls-FLPo (ENS 008, undiluted, 3.6×10^11^ GC/ml), AAV1.ihSyn1.DiO.tTA (Addgene 99121-AAV1, diluted 1:100 to 2.1×10^11^ GC/ml), and AAV1.TRE.fDiO.tdTomato (ENS 068, pEXPR A-183, diluted 1:10 to 2.6×10^12^ GC/ml). The intersectional logic of this three-virus strategy is described in the Results (see ‘Application to sparse and cell-type-specific expression of reporters’).

For validation of cell-type-specific rAAV CRE and FLPo drivers **(Figure 5)**, C57BL/6J mice were injected in the cerebellar cortex, targeting lobule IV/V of the medial vermis (coordinates relative to Bregma: x = 0 mm, y = −6.2 mm, z = −0.5 mm; after a 2-minute wait, the pipette was retracted by 50 µm to −0.4 mm to create a small reservoir at the target), in contrast to the medial DCN target used for **Figure 4**. A total volume of 0.5 µl of diluted virus mixture was injected at a rate of 100 nl/min.

For validation of the c-kit MLI driver **(Figure 5, upper panel)**, mice were injected at P35 with AAV1.hSyn.Flex.GCaMP6f (2.8×10^12^ GC/ml) together with a CRE-dependent rAAV driver carrying a 2.2 kb fragment of the c-kit promoter (Villette et al., 2019), diluted to 7.7×10^12^ GC/ml (upper middle panel) or 7.7×10^10^ GC/ml (upper right panel) to obtain sparse expression, and allowed to express for 10 days before perfusion.

For validation of the pcp2 Purkinje-cell driver **(Figure 5, lower panel)**, mice were injected at P60 with AAV1.eF1α.fDiO.ASAP3b-Kv (1.5×10^13^ GC/ml) together with AAV1.L7_3_.nls-FLPo, an rAAV FLPo-driver carrying a 3 kb fragment of the L7/pcp2 promoter (Nitta et al., 2017), diluted to 2.0×10^12^ GC/ml, and allowed to express for 7 weeks before perfusion.

Each combination shown is representative of 25–30 injections. Histological processing and imaging followed the protocol described below (**Histology and confocal imaging, Figures 4 and 5**).

### Histology and confocal imaging (Figures 4 and 5)

Mice were deeply anaesthetized with an intraperitoneal injection of ketamine (100 mg/kg) and xylazine (10 mg/kg) in saline, then transcardially perfused with phosphate-buffered saline (PBS) followed by 4% paraformaldehyde (PFA) in PBS. Brains were removed and post-fixed overnight in 4% PFA at 4°C. After three washes in PBS, brains were sectioned sagittally on a vibrating microtome (Leica VT1000S, Leica Microsystems, Germany). Sequential 100µ thick sections were collected in PBS and mounted between a glass slide and coverslip using Vectashield Mounting Medium (Vector Laboratories, Newark, CA) or Fluoromount-G (SouthernBiotech, Birmingham, AL). Mosaics and Z-stack images (1 µm optical section thickness) were acquired on an LSM 710 confocal microscope (Zeiss, Jena, Germany) equipped with a Plan-Apochromat 20×/0.8 dry objective (WD 0.55mm). The pinhole was set to 1 Airy unit, and excitation was provided by 488 nm and 561 nm laser lines, with emission collected between 500–530 nm for GFP and 570–620 nm for red fluorescent proteins. Offline image analysis was performed in ImageJ (NIH); several optical slices were combined using the Z-project function to increase the depth of field, and cells were classified visually according to their morphology and spatial location.

### CUBIC tissue clearing and volumetric analysis (Supplemental Figure 4)

For volumetric analysis of axonal terminal density, 3 mm-thick sagittal cerebellar slices were cleared essentially as described in (Matsumoto et al., 2019) and mounted in CUBIC R+. Reagents CUBIC-L, CUBIC-R+, and CUBIC-M were purchased from TCI Biochemicals. Cleared slices were imaged on a Leica SP5 confocal microscope equipped with a 10× XLPLN10XSVMP objective (NA = 0.6, WD = 8.0 mm, Olympus) with CUBIC-M for immersion. Two 3D regions of interest were defined within each imaged lobe: one corresponding to the entire lobe excluding the molecular layer, where no axonal signal was present, and one corresponding to the Purkinje cell layer and superficial granular layer. Mean fluorescence intensity was compared between the two zones. The definition of sections and zones (A and B) and the intensity analysis were done in IMARIS with the 3D quantification tools.

### Cerebellar slice preparation and electrophysiology

For expression of ChR2(H134R)-YFP in cerebellar MLIs **(Figure 7)**, C57BL/6J mice (n = 4) were injected in the cerebellar cortex at the same coordinates and using the same injection parameters as for the c-kit MLI validation cohort **(Figure 5, upper panel**; relative to Bregma: x = 0 mm, y = −6.2 mm, z = −0.5 mm, retracted after a 2-minute wait to −0.4 mm; 0.5 µl injected at 100 nl/min). Mice were co-injected with AAV1.kit.CRE (ENS 001, 4.5×10^11^ GC/ml) and AAV1.DiO.hChR2(H134R)-YFP.WPRE.HGHpA (Addgene 20298, 1.4×10^11^ GC/ml) at P30, and used for acute slice preparation and electrophysiology at D15, as described below.

Acute cerebellar slices were prepared following the procedure described in (Dumontier et al., 2023). Briefly, after deep anesthesia induced with isoflurane (4% in 100% oxygen), mice were decapitated and the cerebellum was rapidly removed and dissected in ice-cold artificial cerebrospinal fluid (ACSF) containing (in mM): 125 NaCl, 3.5 KCl, 1.25 NaH_2_PO_4_, 26 NaHCO_3_, 25 D-glucose, 1.6 CaCl_2_, and 1.5 MgCl_2_, continuously oxygenated with 95% O_2_ / 5% CO_2_. Parasagittal slices (300 µm) were cut using a vibrating blade microtome (Campden Instruments or Leica VT1000-S) in a potassium-rich gluconate cutting solution at 4°C containing (in mM): 130 K-gluconate, 15 KCl, 0.05 EGTA, 20 HEPES, and 25 D-glucose, pH adjusted to 7.4 with NaOH, supplemented with D-APV (50 µM) to prevent glutamate excitotoxicity. Slices were transiently transferred to a mannitol-based recovery solution at 34°C containing (in mM): 225 D-mannitol, 2.5 KCl, 1.25 NaH_2_PO_4_, 25 NaHCO_3_, 25 D-glucose, 0.8 CaCl_2_, and 8 MgCl_2_ (oxygenated with 95% O_2_ / 5% CO_2_, supplemented with D-APV 50 µM), then stored in oxygenated ACSF at 34°C for up to 6 hours before recording.

Electrophysiology recordings were performed in a recording chamber mounted on the AOD-based ULoVE two-photon microscope described below (Villette et al., 2019) perfused at 4 ml/min with oxygenated ACSF at 32–34°C. Slices were visualized using infrared Dodt contrast imaging. Recording pipettes were pulled from borosilicate glass capillaries (outer diameter 1.5 mm, wall thickness 0.225 mm, Hilgenberg).

For Purkinje cell recordings, Purkinje cells were identified by their large soma size and characteristic location in the Purkinje cell layer. Whole-cell voltage-clamp recordings were obtained at a holding potential of −70 mV using 3–5 MΩ pipettes filled with an intracellular solution containing (in mM): 110 CsCl, 20 TEA-Cl, 10 HEPES, 6 NaCl, 10 EGTA, 0.2 CaCl_2_, 4 ATP-Mg, 0.4 GTP-Na, pH adjusted to 7.4 with CsOH. Inhibitory postsynaptic currents (IPSCs) were pharmacologically isolated by adding D-APV (50 µM) and NBQX (2 µM) to the ACSF to block ionotropic glutamate receptors.

For MLI recordings, molecular layer interneurons (basket cells) were identified by their location in the inner molecular layer, small soma diameter (9–10 µm), and ChR2(H134R)-YFP fluorescence. Whole-cell voltage-clamp recordings were obtained using 3–5 MΩ pipettes filled with the same solution as for the Purkinje recordings.

Data were acquired using an EPC10 amplifier (HEKA), sampled at 20 kHz, and filtered at 3–8 kHz. All recordings were performed 15 days after viral injection, consistent with the timeline described in **Figure 7A**.

### In Vivo Optical recordings in mouse cerebellar cortex using ULoVE Animal handling, viral injections, and surgeries

Five wild-type C57Bl/6J adult mice (3 females, 2 males; >60 days old; body weight 20–25 g; n = 2 for GCaMP6f, n = 1 for jRGECO1a, and n = 2 for JEDI2P-Kv) were housed in standard conditions (12-hour light/dark cycles, light on at 07h00., with water and food ad libitum). A preoperative analgesic was administered (buprenorphine, 0.1 mg/kg), and Ketamine-Xylazine was used as an anesthetic (Centravet).

GCaMP6f (Addgene #100835) and jRGECO1a (Addgene #100852) CRE-dependent AAVs were purchased from Addgene; the constructs for the remaining six were generated using the cloning pipeline described above, and the viruses were produced in-house, as described above. For strategy ➀, AAV1.L7_0·6_.CRE was used at final titers of 1×10^12^, 5×10^10^, or 1×10^10^ GC/ml, combined with AAV1.CAG.Flex.GCaMP6f.WPRE.SV40 at 1×10^11^ GC/ml. For strategy ➁, AAV1.L7_0·6_.CRE was used at final titers of 1×10^12^ or 1×10^10^ GC/ml, combined with AAV1.CAG.Flex.NES-jRGECO1a.WPRE.SV40 at 1×10^11^ GC/ml. For strategy ➂, either the combination of AAV1.L7_3_.CRE (5×10^10^ GC/ml) with AAV1.eF1α.Flex.JEDI2P-Kv (5×10^12^ GC/ml) was used, or a dual-intersectional approach combining AAV1.L7_3_.nls-FLPo (2×10^10^ GC/ml) and AAV1.L7_3_.tTA (1×10^11^ GC/ml) with AAV1.TRE.fDiO.JEDI2P-Kv (1×10^12^ GC/ml).

Viruses were combined in a saline solution containing 0.001% Pluronic acid (ThermoFisher 24040032). Volumes of 200–300 nl were injected at a flow rate of 75 nl/min into the cerebellar cortex (lobule V or VIa; coordinates from bregma: anteroposterior 5.8–7.4 mm, mediolateral ±800 µm, dorsoventral –350 µm from brain surface). A custom-designed aluminum headplate was fixed to the skull with layers of dental cement (Metabond). A custom laser-cut #1 coverslip was placed over the cerebellar cortex and secured with dental cement (Tetric Evoflow). Mice were allowed to recover for at least 7 days before recording sessions and were housed in groups of at least 2 mice per cage. Imaging and optical recordings were performed between 8 and 12 days post-surgery. Mice were handled before recording sessions to minimize restraint-associated stress, and experiments were conducted during the light cycle.

### ULoVE optical recordings

Recording sessions lasted 1–2 hours and were conducted while mice behaved spontaneously on top of an unconstrained running wheel in the dark (Villette et al., 2017). Recordings were performed using an acousto-optic deflector (AOD)-based random-access multiphoton system (Karthala System) based on a previously described design (Villette et al., 2019). The excitation was provided by a femtosecond laser (InSight X3, Spectra Physics) mode-locked at 920 nm with a repetition rate of 80 MHz. A water-immersion objective (CFO Apo25XC W1300, 1.1 NA, 2 mm working distance, Nikon) was used for excitation and epifluorescence light collection. The signal was passed through an IR blocking filter (TF1, Thorlabs), split into two channels using a 562 nm dichroic mirror (Semrock), and passed to two H12056P-40 photomultiplier tubes (Hamamatsu) operating in photon-counting mode. The 510/84 filtered green channel was used for collecting GCaMP6f and JEDI2P-Kv signals, while a 607/70 filter was used to collect jRGECO1a red channel signals. Prior to ULoVE optical recordings, the same AOD-based microscope was used in sequential point scanning imaging mode to locate and identify target dendrites. A time series of 50 images at 2 µs/pixel, 0.212 µm/pixel was acquired and post hoc motion registered to produce the images shown in **Figures 6B and E**.

The optimized ULoVE excitation pattern (Lombardini et al., in preparation) reproduced a grid of 3 × 3 points (Villette et al., 2019). This grid spans a 4 × 4 µm area and is scanned diagonally 3 µm in width (x) and 9 µm in height (y), covering a volume of approximately 6 × 15 × 30 µm (x, y, z) to homogeneously fill an extended excitation volume that continually encompasses the dendrite’s cytoplasm and plasma membrane. To obtain a good signal-to-noise ratio, two excitation patterns were used per acquisition window of 80 µs, yielding a temporal resolution of ∼ 5 kHz. For imaging, laser power was set to deliver 20 mW post-objective and pre-sample, then adjusted for mono-exponential loss through tissue with a length constant of 170 µm. For ULoVE optical recordings, power was multiplied by 2.5 relative to the imaging mode value to account for the greater excitation volume. The applied power never exceeded 200 mW. Dendrites were selected on the basis of being sufficiently bright to obtain a large signal-to-noise ratio while being spatially isolated to avoid contamination from neighboring structures. Recordings were stopped after 3 minutes.

For each dendrite recording, two ULoVE volumes were simultaneously used as regions of interest (**Figure 6C**); fluorescence traces therefore represent the sum of photons collected from both volumes. No obvious photobleaching was observed except in a minority of jRGECO1a recordings and in most JEDI2P-Kv recordings. For the latter, a moving median computed over a 100 ms time window was used to subtract the drifting baseline. Baseline fluorescence (F_0_) was estimated as the mode of the raw photon-count distribution (200-bin histogram), and ΔF/F (%) traces were computed as (F − F_0_)/F_0_ × 100. Voltage ΔF/F (%) traces were computed with inverted sign relative to calcium, as (F_0_ − F)/F_0_ × 100, to reflect the polarity of the JEDI2P-Kv indicator. Note that smoothing applied for display purposes (3 ms Gaussian kernel for calcium traces, 0.5 ms Gaussian kernel for voltage traces; see **Figure 6D** and **F**) differs from the kernels used in the analyses described below.

### Event detection

#### Calcium transients

For GCaMP6f and jRGECO1a recordings, candidate events were identified from a delayed derivative computed on a lightly smoothed fluorescence trace (1 ms Gaussian kernel), subtracting a peak segment (15 ms) from a baseline segment (50 ms) separated by a 20 ms gap at each time point of the 5 kHz optical recording. Local extrema of the Z-scored delayed derivative trace were detected with a peak-finding algorithm using a minimum peak height and prominence of 1.1 and 1.0, respectively, and a minimum inter-event interval of 30 ms. Candidate events were further screened using the ratio of post-to pre-event mean amplitude in a peri-event raster, with a data-driven threshold (density-based, validated by visual inspection) separating true transients from noise fluctuations.

#### Voltage transients

For JEDI2P-Kv recordings, the same delayed derivative detection was applied with shorter rise duration (3 ms) and peak (8 ms) windows, appropriate for the faster kinetics of the voltage indicator, and a fixed Z-score threshold (height/prominence = 5, minimum inter-event interval 30 ms). Detected onsets were realigned by cross-correlation of individual event waveforms to a reference, yielding a per-event temporal shift used to correct event times. Events whose pre-onset baseline (raster window −15 to −2 ms) deviated from the population distribution were excluded as putative false positives.

### Kinetics characterization and voltage event waveform analysis

For each recording, isolated events (no neighboring detection within the analysis window) were selected from peri-event rasters built on the smoothed ΔF/F trace. Rasters were baseline-normalized to a short pre-event window (−1.2 to 0 ms). Rise and decay kinetics (time constants τ_rise_ and τ_decay_) were obtained by fitting the averaged event waveform with an exponential rise/decay model. Distributions of τ_rise_ and τ_decay_ were pooled across calcium indicators and fitted with a Gaussian model to characterize population kinetics; the resulting calcium rise-time distribution was compared to the voltage-indicator τ_rise_ distribution obtained from the dendritic population raster analysis.

For the fast voltage signals, individual normalized event waveforms were further screened with a peak-detection algorithm applied to a lightly smoothed trace (0.5 ms Gaussian kernel) to identify multi-peak events, classified as doublets or triplets based on their number of peaks. The inter-peak interval distribution within multi-peak events was fitted with a double-Gaussian model to characterize the underlying bimodal timing structure. Inter-peak intervals were also used to separate closed and separated two-peak events: for the closed category, the first peak occurred between 0.6 and 1.3 ms after onset and the second peak between 3.8 and 4.6 ms; for the separated category, the second peak occurred between 5 and 10 ms. Selection criteria for three-peak events were: first peak between 0.6 and 1.3 ms, second peak between 3.8 and 4.6 ms, and third peak between 8 and 10 ms. The time-frequency content of each event category was characterized using a continuous Morlet wavelet transform, with dominant oscillatory frequencies identified as peaks in the power-to-frequency profiles. AHP amplitude was measured as the difference between the mean trough (15–55 ms following onset) and the local baseline (mean of the 15 ms window prior to onset).

All analyses were performed in MATLAB 2019b (MathWorks). Code will be made available, and data can be shared upon request.

### Two-photon photostimulation of MLIs using ULoVE

Two-photon photostimulation of ChR2(H134R)-YFP-expressing MLIs was performed using the AOD-based ULoVE microscope described above. The excitation wavelength was set to 920 nm. Rather than the extended volume excitation pattern used for in vivo calcium and voltage recordings, photostimulation of individual MLI somata was performed using a doughnut-shaped excitation pattern generated by scanning a diffraction-limited focus along a circular trajectory using the acousto-optic deflectors (Villette et al., 2019). A series of 20 doughnuts was designed to match the typical MLI soma diameter of 9–10 µm, with each doughnut scanned for 70 µs, yielding a total single-cell excitation duration of 1.4 ms per photostimulation cycle.

For characterization of photocurrent input–output relationships (**Figure 7D**), laser power was varied to span squared powers of 11.3, 42.5, 149 and 215 mW^2^, and photocurrents were recorded in whole-cell voltage clamp from individual ChR2(H134R)-YFP-expressing MLIs under concurrent photostimulation. For spatial resolution measurements (**Figure 7E**), photocurrent amplitude was measured as a function of lateral and axial displacement of the ULoVE focus relative to the center of the MLI soma, with the laser power held constant.

For multi-cell sequential photostimulation and functional connectivity mapping (**Figure 7F, G**), nine MLIs in the vicinity of a simultaneously patch-clamped Purkinje cell were targeted in sequence at a revisit cycle frequency of 710 Hz, with each MLI receiving one 1.4 ms multi-doughnut excitation per cycle. Purkinje cell IPSCs were recorded in voltage clamp at −70 mV during sequential MLI photostimulation. Ten trials were acquired per cell to assess the reliability of any detected synaptic connections.

## Supporting information

Supplemental_AAV_production_worksheet

Gateway_ULoVE_Figures_2,_3_&_4 Supplemental_v3_bioRxiv

## Acknowledgments

This work was supported by grants from Région Île-de-France to the qPCR-HD-Genomic Paris Centre Core Facility, ANR grants: PEPR, (ANR-22-PEBI-0012); MuVriCC, (ANR-20-CE16-0018); FucPoc, (<u>ANR-20-CE16-0026)</u>; ANR-24-INBS-0005 FBI BIOGEN; ANR-10-LABX-54 MEMOLIFE & OptoConnectics, (ANR-25-CE16-7612), NIH BRAIN grants: <u>U01NS133971</u> & <u>U01NS103464</u>. Confocal Imaging was done in the imaging facility SCM, INSERM US36 | CNRS UAR2009 | Université Paris Cité (<u>Biomedtech-facilities</u>), and at the IBENS imaging facility, a member of the National Infrastructure France-BioImaging, and supported by the DIM C-BRAINS, funded by the Conseil Régional d’Île-de-France de France(<u>IMACHEM-IBiSA).</u>

Virus titering was done at the ENS High-Throughput qPCR core facility; http://www.qpcr.cnrs.fr/.

We thank Mélissa Desrosiers and Marine Delagrange for technical assistance and Nathalie Pardigon for critical reading of the manuscript.

During the preparation of this manuscript and its associated figures, the authors used Claude (Anthropic) to assist with data visualization, figure generation, and manuscript text editing. The authors reviewed, verified, and take full responsibility for all content.

## Author Contributions

A.J., C.E., A.T., & B.M. performed the CUBIC imaging, and C.E. did the analysis; A-P. B. provided reagents; I.L., S.D. & J.B. conceived of the project; A.A. and J.B. performed the cloning; J.B. produced the viruses with help from A.A. and C.M.-H.; A.J. did the histology and confocal imaging; J.B. and V.V. performed the animal surgeries: V.V. did the in vivo experiments and analysis; J.B. wrote the manuscript with input from V.V. and L.B.

## Notes

### Competing Interest Statement

The authors have declared no competing interest.

### Summary of Updates

Version 3. This version replaces v2, in which the figures were reduced in resolution during conversion from a Word file; all figures are now provided at higher resolution in a PDF. The text and figures were also revised in response to peer review: (1) specified the limitations of restriction-enzyme cloning in the Introduction; (2) clarified in the Results, Figure 6 legend and panel 6A that each indicator strategy was delivered to separate animals and that calcium and voltage were not recorded simultaneously; (3) corrected statements about the source of constructs (GCaMP6f and jRGECO1a were obtained from Addgene; the Gateway-built components are the driver viruses and the JEDI2P-Kv vectors); (4) revised Discussion statements about prior literature on dendritic spikelet recording; (5) corrected terminology and abbreviations (titer; PV, SST, VIP, CCK, GAD67; DCN); (6) updated Figures 3, 4, 5, 6A and 7A and Supplements to Figures 2, 3 and 4 (enlarged text throughout, simplified Figure 4 schematic, updated promoter nomenclature: hSyn, GfaABC1D, c-kit); (7) added two supplementary spreadsheets, the AAV production worksheet and Supplemental Table 1 (production record of the 105 AAV productions), and a header page to the supplementary figures file.

