## Supplementary material for "Streamlined Production of Recombinant Adeno-Associated Viruses Using Gateway Cloning Technology for Neural Circuit All-Optical Interrogation": Gateway_ULoVE_Figures_2,_3_&_4 Supplemental_v3_bioRxiv

**This file contains the supplementary figures cited in the main text. Supplement to Figure 2 shows how the entry clones are made: BP recombination of a PCR-amplified promoter (A) and Gibson assembly of a transgene into a pre-configured Flex backbone (B). Supplement to Figure 3 is a Sankey diagram of the 69 in-house rAAV expression constructs, tracing each through its capsid, promoter, recombinase switch, gene of interest, trafficking motif and post-transcriptional element; the full indexed production record is in Supplemental Table 1 (Excel file, provided separately). Supplement to Figure 4 shows anterograde tracing of vGlut2+ DCN axons into the cerebellar cortex in CUBIC-cleared tissue. The worksheet used to plan the transfections and to follow the harvest protocol is provided as a separate Excel file (Supplemental AAV production worksheet).**

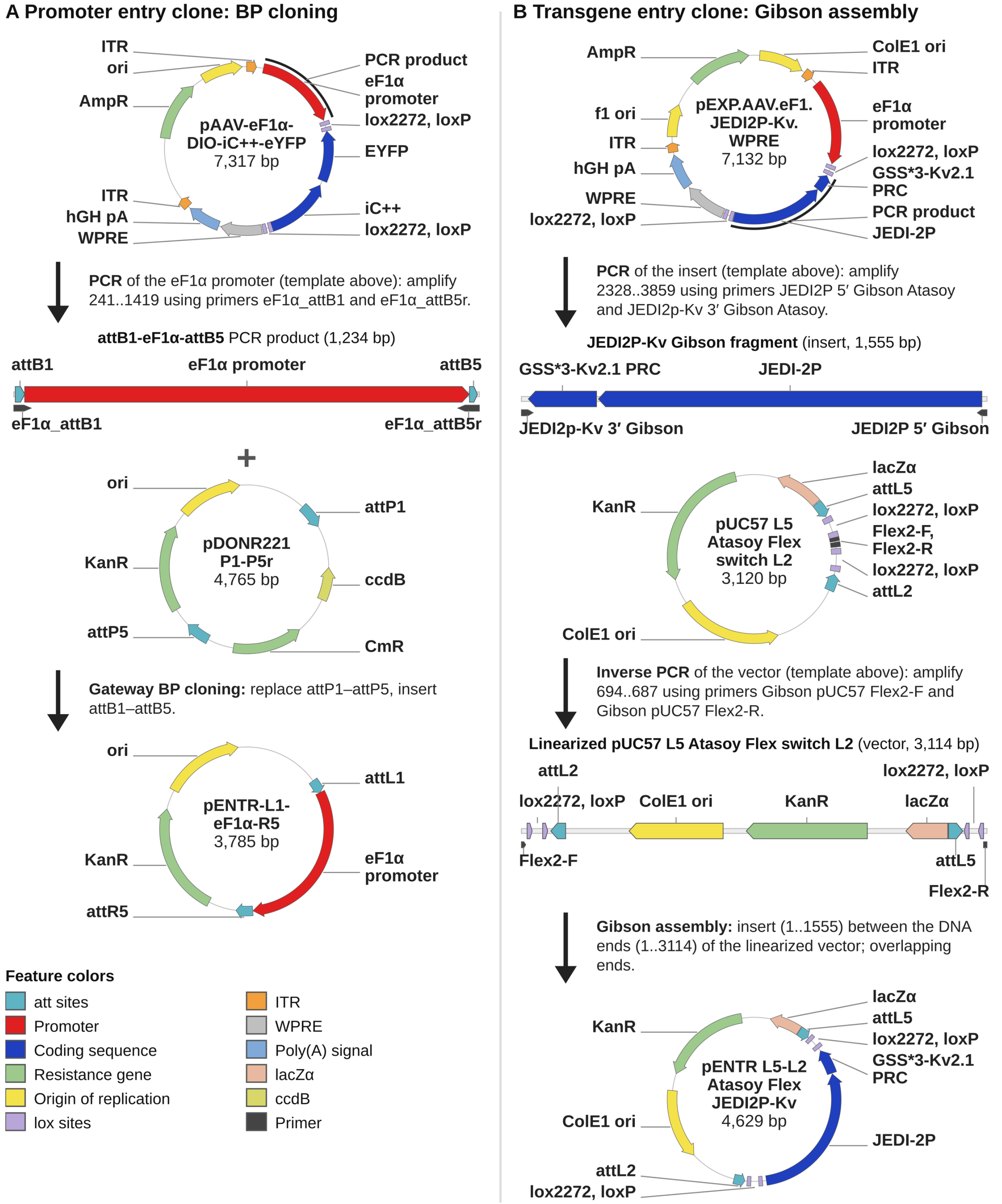


**Supplement to Figure 2. Generation of pENTR plasmids by BP recombination and Gibson assembly.** **(A)** The eF1α promoter was amplified from pAAV-eF1α-DIO-iC++-eYFP with primers appending attB1 and attB5r sites and recombined by BP Clonase with the attP1 and attP5 sites of pDONR221 P1-P5r, excising the ccdB/CmR cassette to give pENTR-L1-eF1α-R5 (promoter flanked by attL1 and attR5). KanR selection and ccdB counter-selection ensure correct recombinants. **(B)** For conditional expression, backbones carry heterotypic lox (Flex) or frt (fDiO) sites (Atasoy et al., 2008; Fenno et al., 2014). JEDI2P-Kv (1,555 bp) was amplified from pEXP.AAV.eF1.JEDI2P-Kv.WPRE with ends overlapping the insertion site of the vector, which was linearized by inverse PCR of pUC57 L5 Atasoy Flex switch L2 (3,114 bp). Gibson assembly inserted JEDI2P-Kv in the anti-sense orientation between the lox sites, giving pENTR L5-L2 Atasoy Flex JEDI2P-Kv (4,629 bp). Colors indicate feature types; black arcs mark amplified regions.

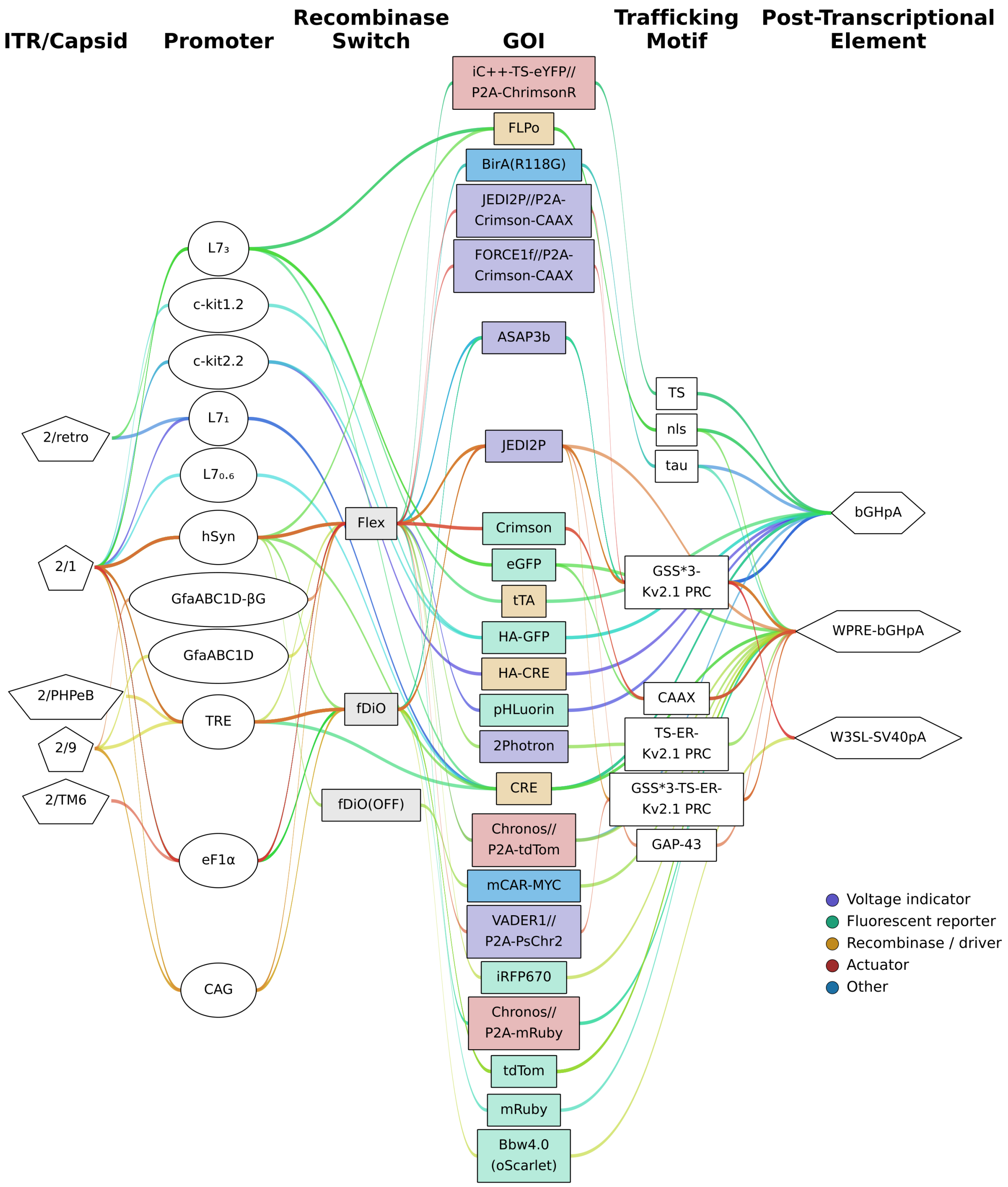


**Supplement to Figure 3. Sankey diagram: Diversity of the 69 different rAAV expression constructs produced using the Gateway cloning pipeline.**

Each colored line represents one Gateway construct produced with the cloning pipeline, tracing the component elements of that construct from left to right in canonical expression order: ITR/capsid types, promoter, conditional expression configuration, GoI (GEVI/reporter/recombinase or transactivator/actuator or other), subcellular targeting motif, and post-transcriptional regulatory elements.

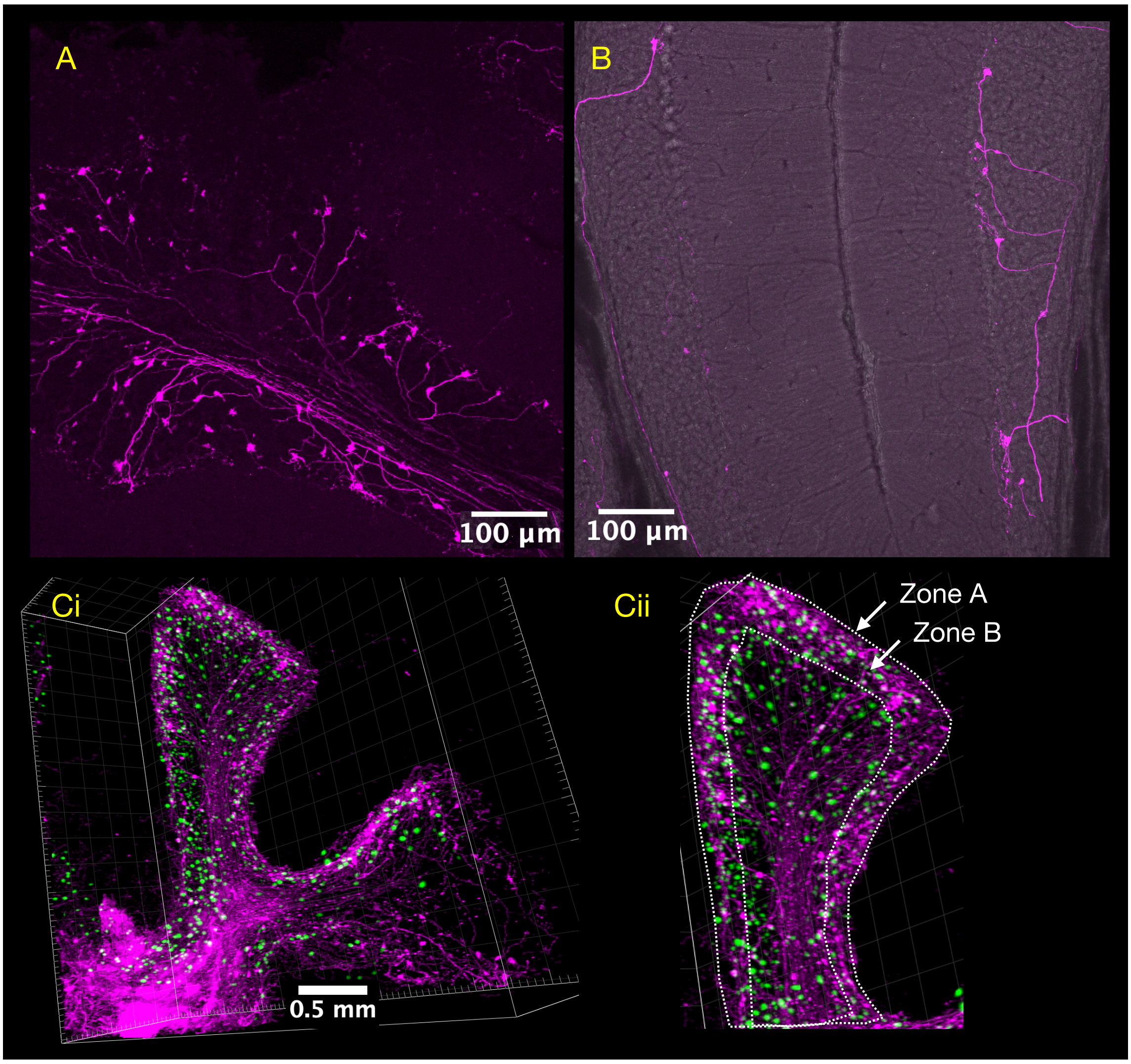


**Supplement to Figure 4. Anterograde tracing of vGlut2+ DCN axonal projections to the cerebellar cortex. (A)** Confocal image of a vibratome section showing tdTomato-labelled axons (magenta) from vGlut2+ DCN neurons, labelled using the triple-intersectional strategy described in **Figure 4**, with dense terminal boutons concentrated at the boundary between the granule cell layer and the molecular layer **(see Figure 5, upper left panel, for a schematic of the cerebellar cortical cytoarchitecture)**. Scale bar, 100 µm. **(B)** Confocal image from a separate section with sparser density of fibers, showing individual tdTomato-labelled vGlut2$^{+}$ DCN axons (magenta) and their terminal boutons at the granule cell layer–molecular layer boundary. Scale bar, 100 µm. **(Ci)** 3D reconstruction of a cerebellar lobe imaged in a 3 mm-thick sagittal slice, CUBIC-cleared and mounted in CUBIC R+ (Leica SP5, 10× XLPLN10XSVMP objective (NA = 0.6, WD = 8.0 mm, Olympus), showing tdTomato-labelled DCN axons and terminals (magenta) and GlyT2::eGFP-labelled glycinergic neurons (green, as in **Figure 4**). Scale bar, 0.5 mm. **(Cii)** Z-projection of the same reconstruction with two 3D zones outlined for quantitative comparison: Zone A, corresponding to the entire lobe excluding the molecular layer (where no axonal signal is present), and Zone B, corresponding to the Purkinje cell layer and superficial granular layer. Mean signal intensity in Zone B was 40% higher than in Zone A (Zone A: volume $2.16 \times 10^{8}$ µm$^{3}$, mean intensity 0.589; Zone B: volume $1.93 \times 10^{7}$ µm$^{3}$, mean intensity 0.829), consistent with the preferential targeting of DCN axon terminals to the granule cell layer–molecular layer boundary observed in **A** and **B**.
